# Neural competition and probabilistic representations

**DOI:** 10.64898/2026.08.31.748371

**Authors:** Nava Rubin, Jorge Lobo

**Affiliations:** Institució Catalana de Recerca i Estudis Avançats, Barcelona 08010, Spain; School of Engineering, Universitat Pompeu Fabra, Barcelona, 08018, Spain

**Author notes:** Deceased (Nava Rubin).

**Keywords:** Balanced networks, competition, attractors

## Abstract

Perception and action show tight links to the statistical structure of physical stimuli and likely rewards, but the underlying mechanisms are unknown. A simple, biologically plausible network model shows that probabilistic behavior emerges naturally in diverse scenarios, and arises from sampling of competing responses. Notably, it also provides a principled computational rationale for the prevalent finding of balanced excitation and inhibition in the brain. Recurrent connections within a population of excitatory neurons embed multiple attractor states, and coupling to a pool of inhibitory neurons enforces mutual exclusivity among the states. Upon concurrent stimulation, competing attractors alternate in activity. These global state transitions are caused by local, uncorrelated spiking noise and yet convey, over time, the relative strengths of attractors’ support. The simplest *probabilistic competitive recurrent networks* (PCRNs) allow for closed-form analysis, shedding light on the neural basis of choice behavior under uncertainty. More complex systems of laterally connected PCRNs can collectively resolve the myriad local ambiguities pervasive in sensory stimuli, rapidly settling into globally-consistent configurations that match perceptual reports. Alternations are crucial in all cases, and occur only if a PCRN’s inhibitory pool is strong enough to prevent attractors’ activity from reaching saturation. A balance between excitation and inhibition is thus both a prerequisite and a hallmark of probabilistic sampling in cortical networks.

## 1 Introduction

The scenario addressed by the model concerns the need to choose between different possible responses in the face of conflicting or ambiguous inputs. Here ‘input’ refers to stimulation from sources external to the network, which may be sensory, intra-cortical, or both, and a ‘response’ is realized by convergence to one of the network’s attractor states. (Output connections are not assumed but can be added as needed.) Each attractor state is a different stable pattern of activity, embedded via Hebbian connectivity among the excitatory neurons. A basic tenet of the model is that the patterns embedded within its excitatory population correspond to different options which are mutually exclusive by their very nature – eg, they may represent different edge orientations in the same location in space, or code for a limb’s movement in different directions. In this context, there is an obvious challenge posed by external inputs that stimulate more than one of the patterns, as they may lead to concurrent activation of contradictory percepts or actions. Preventing such undesirable outcomes is a major role of the PCRN’s inhibitory population. It delivers a uniform negative feedback which regulates the overall activity in the network, allowing for only one of the embedded patterns to be active at any given time. Choice is thus achieved via *neural competition*, whereby excitatory cell assemblies representing the different options are forced to compete for activity (Fig. 1(a)). The use of inhibition to enforce mutual exclusivity (XOR) is common in models of cognitive and perceptual choice (Laing and Chow, 2002; Wang, 2002), but the present model differs in an important way. Previously, there has been a (mostly tacit) assumption that achieving XOR is tantamount to obtaining ‘winner takes all’ behavior, but the two concepts can be dissociated (Hahnloser et al., 2000). In the current model, the inhibitory feedback not only enforces XOR between the competing attractors, but also serves to regulate the activity of the ‘winner’ so that rather than taking ‘all’, it only reaches a level proportional to its input strength. This creates the possibility for ‘apportioned winning’ – over time, or trials – which is realized in a PCRN via spiking noise.

**Figure 1:**
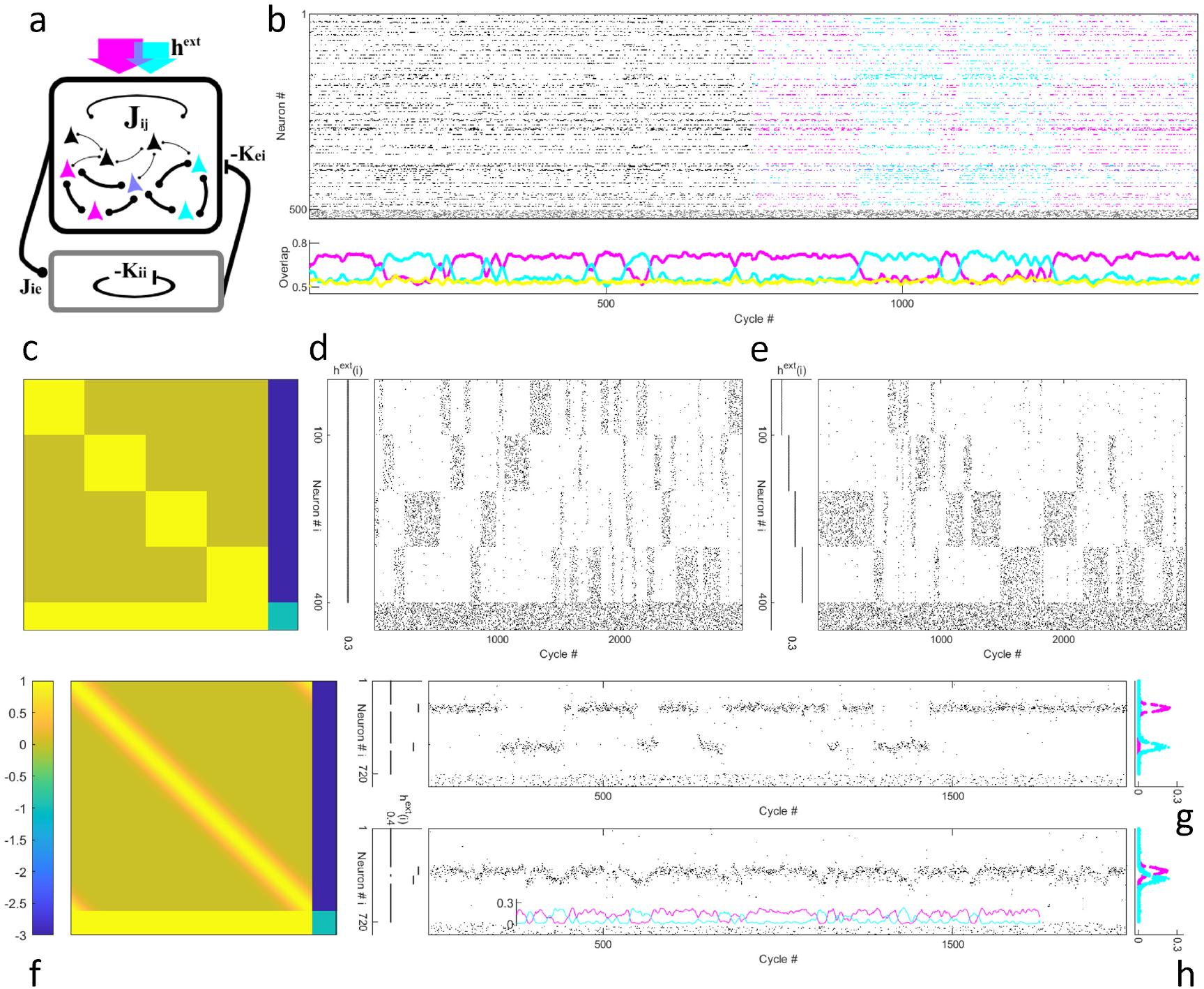
Neural competition leads to probabilistic sampling in diverse scenarios. **a.** The PCRN model architecture. An excitatory neuronal population is interconnected by Hebbian synapses (top box), and coupled to a common pool of inhibitory neurons (bottom) via reciprocal connections of uniform strength. External stimulation (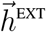, top arrows) may simultaneously potentiate two or more excitatory cell assemblies (magenta and cyan), possibly with overlap (lilac). **b-h**. Simulations of activity in three example PCRNs, each embedding a different set of patterns, in response to time-constant, noise-free 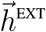. **b**. Ten random binary patterns were embedded in an excitatory population of 200 neurons, and 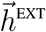 targeted the neurons belonging to two of the patterns, with unequal strength. Sampling behavior is hard to discern from the raster (top) when neurons are unlabeled (cycles 1-750, monochrome), but clear in the labeled portion (cycles 751-1500) and in time-courses of the overlap between network state and each pattern (bottom; yellow shows an unstimulated pattern). **c,f**. Connections within the excitatory population comprise a symmetric sub-matrix (Eq.1), flanked by three blocks of uniform values that render the full connectivity matrix asymmetric (Eq.2). **c-e**. With four binary patterns of contiguous, non-overlapping sites, sampling appears in the rasters as intermittent blocks of activity and quiescence. Under uniform stimulation (panel d, flat black line), each of the competing sub-populations is active a quarter of the time, on average. When *h*^EXT^ values differ between populations, the network spends more time in those more strongly stimulated (panel e). **f-h**. Sixty densely overlapping Gaussian patterns, embedded in a PCRN of 600 excitatory neurons, produce a smooth diagonal ridge in the connectivity matrix (panel f). Stimulation at two well-separated loci results in visible alternations between distinct neuronal groups (panel g), but states remain differentiated even with minimally-separated inputs (h; magenta and cyan time-courses show mean activity in neuronal subsets stimulated by higher and lower peaks, respectively). In both cases, neurons’ activity profiles (right-most blocks, panels g and h)) do not mirror the boxcar shape of external stimulation (black lines, left), but rather follow the embedded patterns, indicating a dynamics governed by attractor states. Here and in later figures *J*_IE_=*K*_II_=1, *K*_EI_=3.

Humans and animals exhibit variability in their choice behavior, even when relevant environmental factors remain unchanged. This variability is thought to arise from neural noise, but how and why it comes about is not well understood. Positing that the neurons computing the response have a shared noise source is problematic because peripheral sensory neurons show highly regular firing and, within cortex, there are no known mechanisms for correlated fluctuations to generate a putative ‘background’ noise. By contrast, synaptic processes such as ion channel gating and quantal release are well established sources of neural noise, so that spiking is better described probabilistically, as a smooth (if steep) sigmoid function of the input, rather than a step function (Calvin and Stevens, 1968; White et al., 2000). Since spiking noise occurs independently in each neuron (‘local’), it would average out of the mean firing rate of large neural populations, and may therefore appear unsuited to account for variability observed at the behavioral level (‘global’). But the analysis and simulations of PCRNs presented here suggest otherwise.

In all that follows, spiking noise is the only source of variability: external stimulations will be kept constant, and no other inputs are posited. And yet, a distinct form of global variability emerges: the identity of the active pattern is not fixed, nor is it determined solely by which gets the stronger stimulation. Instead, the competing attractors alternate stochastically, each getting to win a fraction of time reflecting its relative strength (Figs. 1b-h). Alternations proceed at a mean rate that depends on single neurons’ level of spiking noise, and can be made arbitrarily slow (or fast) by adjusting the slope of the sigmoid probability function of spiking. This provides a general mechanism of probabilistic sampling that can operate across a range of time-scales in a variety of circumstances.

## 2 Model formulation

The PCRN neurons are binary elements which can be either active (‘firing’, *V*_*i*_ = 1) or quiescent (*V*_*i*_ = 0). For a population of N_E_ excitatory neurons, the embedded patterns are a set of *P* vectors 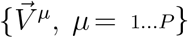 of length N_E_ each, with elements representing the prescribed firing rate of the corresponding neuron. The connectivity within the excitatory population is given by the N_E_ *×*N_E_ symmetric matrix

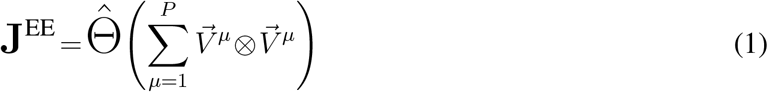

where 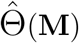 is a monotonic operator that transforms the elements of **M** to be of O(1). Note that the patterns must be non-negative, but they may be either binary (as in Fig. 1(c)) or continuous variables (as in Fig.1(f)). Accordingly, 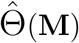 is set as the element-wise step function for binary patterns, and a division by *max* ({M_*ij*_}) for continuous patterns.

Alongside a clear similarity with Hopfield’s (Hopfield, 1982) formulation of a Hebbian connectivity matrix, an important difference to note is that the latter consisted of both positive and negative elements. In contrast, **J**^EE^ prescribes all-positive connectivity which, if left unchecked, would result in activity spreading throughout the population for all but the simplest cases of patterns with little to no overlap. What prevents such undesirable outcomes is the PCRN’s inhibitory population, which provides negative feedback and regulates the overall activity in the network. A significant advantage of this architecture is that activity regulation can be achieved with the strengths of connections to-, from- and within the inhibitory population taken as uniform. This means that just three additional parameters are needed to fully specify connectivity in the PCRN, which greatly simplifies its analysis. Denoting by *J*_*xy*_ and *−K*_*xz*_ the strength of connections made onto neuron *x* by an excitatory and an inhibitory neuron, respectively, the three parameters are the positive scalars *J*_*ie*_, *K*_*ei*_ and *K*_*ii*_. With an inhibitory population of size N_I_, the full network connectivity is thus given by the matrix

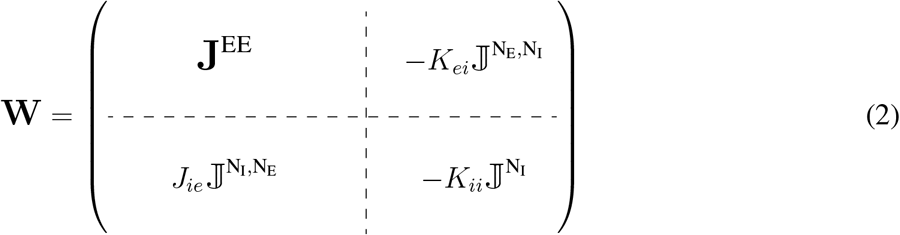

with J^*m,n*^ and J^*m*^ denoting the all-ones matrices of sizes *m×n* and *m×m*, respectively (for examples see Fig. 1, panels (c,f)). The total input experienced by neuron *i* corresponds to the *i*-th element in the vector 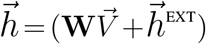, where 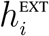 denotes the external stimulation to it. Neurons update their state, independently and asynchronously, with the probability of spiking set to

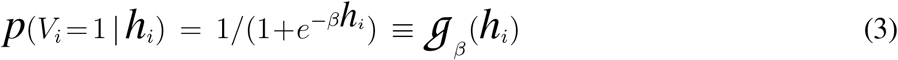

The dynamic mean-field equations describing the system (Glauber, 1963) are then given by

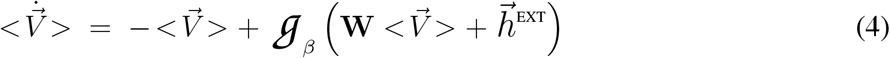

(Activation thresholds were set to 0; for any other value, 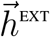 can be shifted accordingly.) The parameter *β* controls the level of spiking noise in the network: by raising the slope of the sigmoid function *G*, it increases the faithfulness with which neurons become active upon positive stimulation (and quiescent for negative input). At the limit *β→∞* the sigmoid turns into a step function and neurons behave deterministically. But for any finite *β*, however large, activity can be predicted only on average: a neuron always has a finite chance of a response opposite to that prescribed by its input. This is akin to thermal noise in physical systems, with 1*/β* being formally equivalent to temperature in the dynamics of dipole-spin interactions (Glauber, 1963; Hinton and Sejnowski, 1983). The term ‘temperature’ will thus be used here as shorthand, to refer to the specific noise mechanism of probabilistic spiking given by *G*_*β*_ (though, note, it need not be thermal in origin). By the same token, when the neurons are deterministic the system is said to be at ‘zero temperature’.

The case of P binary, non-overlapping patterns will be of particular interest. The excitatory neurons then divide into P sub-populations, each having uniform internal connectivity and no connections to the other P*−*1 groups (Eq. 1), except indirectly via the inhibitory population (Eq. 2). Assuming (with no loss of generality) that each group is in contiguous locations in 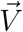 gives the **J**^EE^ matrix a P*×*P block-diagonal structure, underscoring the role of the common inhibitory pool as the sole mediator of influence between the P excitatory blocks (eg, Fig. 1(c)). Denoting the sub-populations by E^*µ*^ and setting the external stimulation to each at a uniform strength 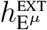, all 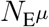 neurons in E^*µ*^ are then identical in terms of their mean-field behavior. This significantly reduces the dimensionality of the problem, with the system of 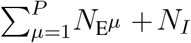 equations in Eq. 4 becoming just P+1 scalar equations, Eqs.5, where *<V*_*x*_*>* denotes the expected value of a neuron in population *x*∈ {E^1^,…, E^P^, *I*}.

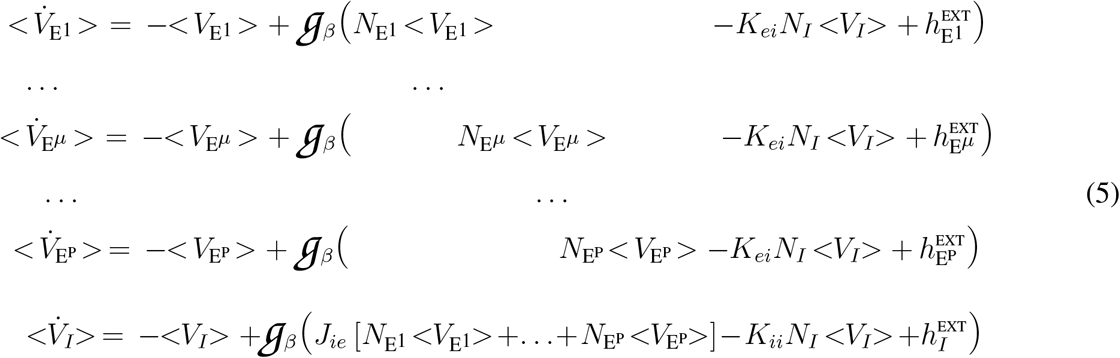

The simplicity of Eqs.5 lends them to formal analysis which will also shed light on the model’s behavior in more complex cases, of competition between attractor states prescribed by overlapping and/or continuous-value patterns. Moreover, for P = 2 the analysis explains key findings from perceptual bi-stability, including several longstanding observations that have been, thus far, difficult to reconcile.

## 3 Zero temperature analysis

Generally, the nonlinear differential equations in Eq. 5 can be solved only numerically. But in the special case of deterministic neurons, the sigmoid *G* becomes a step-function, and closed-form solutions can be derived. Those solutions will prove invaluable, in turn, for understanding what is ultimately of interest here – the behavior of PCRNs in the presence of spiking noise (‘finite temperature’). The analysis given below could be applied to any set of connectivity parameters, but is facilitated by choosing a particularly simple set, for which the model still exhibits the full repertoire of behaviors of interest.

Setting *J*_*ie*_ =*K*_*ii*_ = 1, we can drop the subscripts from the remaining connectivity parameter and define *K*_*ei*_*≡K*. Focusing on scenarios where the inhibitory population is comprised of ‘interneurons’, we set 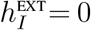. Lastly, assuming that *N*_E*µ*_ is equal to *N*_EP_ *≡ N*_*E*_*/P* for all *µ*, we can factor it out by defining *r*_I|E_ *≡N*_*I*_*/N*_EP_ and 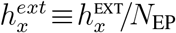. For fixed-point solutions 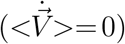, Eqs.5 thus become

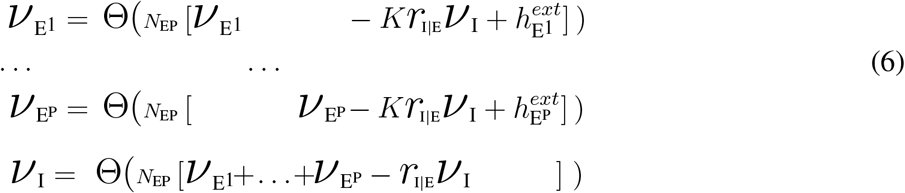

where *νx* denotes the value of *<V*_*x*_*>* at the fixed-point for a neuron in subpopulation *x* (ie, its steady-state firing rate), and Θ(*h*) *≡G* _*β→∞*_ (*h*).

The solutions to Eqs. 6 can be found by testing, case-by-case, all sets of permissible values for the P + 1 unknowns. Since Θ(*h*) assumes the value 1 for all positive inputs and 0 for all negative inputs, naturally the first set of fixed-points to consider are those where *V*_*x*_ is either 0 or 1 for all *x∈* {E^1^,…, E^P^, I}. There are 2^*P* +1^ such possible configurations. For any one of them to be a valid fixedpoint solution, the resultant arguments to Θ must be self-consistent with the posited value of *V*_*x*_ for each of the subpopulations. The given configuration may turn out to be a self-consistent solution in some portion of parameter space (ie, for some ranges of 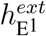, …, 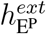 and *r*_I|E_), in the entire space, or not at all. For small P it is straightforward to test all possible configurations, and the results will be instructive also about large P.

Consider first P=1, a single excitatory population with uniform internal connectivity. It is readily verifiable that the pair [*V*_E_, *V*_*I*_]=[0, 0] renders Eqs.6 self-consistent only for 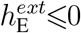, ie, in the absence of external stimulation (here E replaces E^1^ to denote the excitatory population). Conversely, for the pair [*V*_E_, *V*_*I*_]=[1, 1] to be a valid fixed-point the external stimulation must satisfy 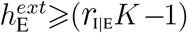, and *r*_I|E_ must be ⩽1. It is also easy to work out that the remaining configurations, [0, 1] and [1, 0], cannot be self-consistent as fixed-points for any value of the external input. The case of P=2 is slightly more cumbersome yet, essentially, yields similar results. With the subscripts A and B replacing E^1^ and E^2^, case-by-case testing reveals that the triplet [*V*_*A*_, *V*_*B*_, *V*_*I*_]=[0, 0, 0] is valid as a fixed-point only when both 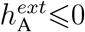 and 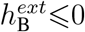; that for one or more of the triplets [1, 1, 1], [1, 0, 1], [0, 1, 1] to be a solution of Eqs. (6), the external input must satisfy 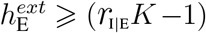 for at least one *E ∈* {*A, B*}; and that the remaining four of the 2^(2+1)^ possible triplets are not self-consistent in any portion of parameter space. What happens when the external inputs are positive, yet weaker than *r*_I|E_*K−*1 ? In this intermediate regime, an altogether different kind of fixed-points comes into play. Note first that although the function *G*_*β*_ (*h*) assumes the value 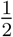 at *h*=0 for any finite value of *β*, at zero temperature (ie, at the limit *β → ∞*) its value is undefined. This means there could be fixed-points where *V*_*m*_ assumes an *intermediate* value for one or more sub-populations *m ∈*{E^1^,…, E^P^, I}. For the full set {*V*_*x*_} to be a self-consistent solution requires that it produces a null argument for Θ in the row(s) corresponding to *V*_*m*_ in Eqs.(6). For the simple case of P=1 there is just one such fixed-point, given by

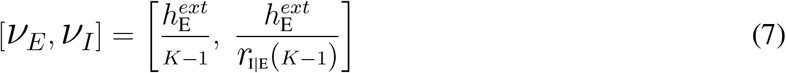

and valid throughout the intermediate range where no solutions with *V*_*x*_=1 or = 0 exist. Networks of P*≥* 2 excitatory sub-populations have analogous fixed-points, but with only one of them active at the intermediate-*ν* level (along with the inhibitory population). In particular, if 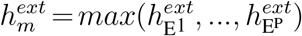 then the vector

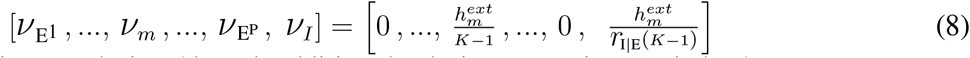

is always a self-consistent solution (though additional solutions co-exist; see below).

Simulation results confirm that the model indeed converges to the intermediate-*ν* fixed-points predicted by Eqs. 7 and 8 when stimulations weaker than *r*_I|E_*K−*1 are applied (Fig. 2). For P = 1, the fractions of active neurons in both the excitatory and inhibitory populations are proportional to 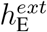, and remains unchanged as long as the latter is constant (panel a). In a P=5 network, the excitatory population receiving the strongest external stimulation is active while the others are all quiescent (panel b). Here, too, the fraction of active neurons is proportional to the external input, but now it is the maximal input that counts, regardless of stimulation values to the other sub-populations. The P*≥*2 network thus offers (at zero temperature) a novel solution to a well-known challenge in neural computation, that of simultaneously performing digital selection and analogue amplification (Hahnloser et al., 2000), also known as the *MAX* operation (Yu et al., 2002).

**Figure 2:**
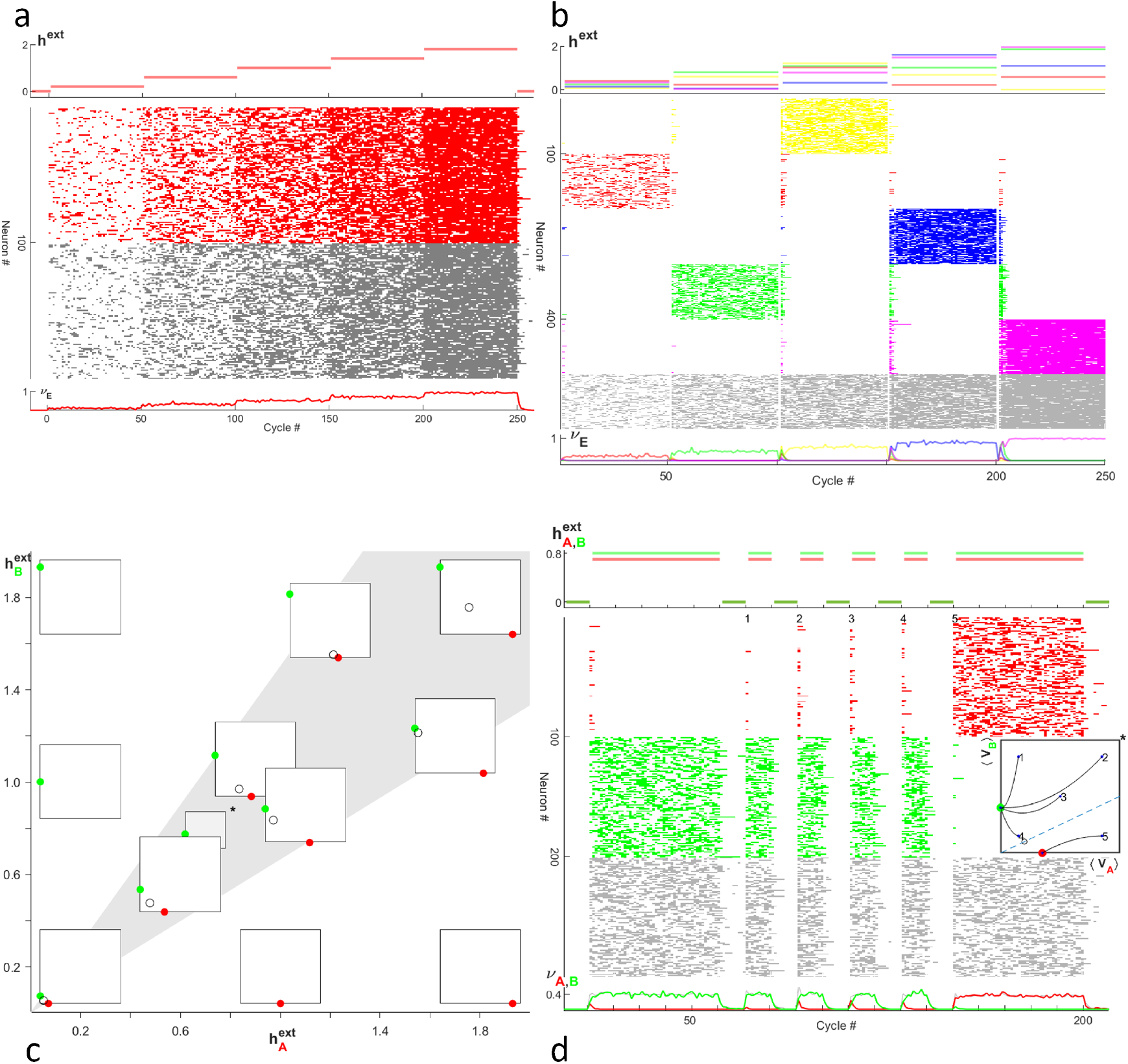
Deterministic neurons in uniformly connected sub-populations. **a-b**, d.Simulation results from networks embedding *P* binary non-overlapping patterns, for *P*=1, 2 and 5 (*N*_EP_=*N*_*I*_=100 in all cases). Neurons were updated asynchronously in pseudo-random order. The top graph in each panel shows 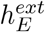 over time, rasters show the resultant activity, and the bottom curves show mean activity for each sub-population. **a**. Neurons in *E* received uniform stimulation in five segments of 50 cycles each, stepped up from 0.2 to 1.8. Mean activity in the *E* and *I* populations increases linearly with 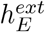, closely matching *V*_*E*_ and *V*_*I*_ predicted by Eq. 7. **b**. Rising steps of constant stimulation were applied, in separate segments, to one (randomly picked) of *P* =5 sub-populations, while all remaining received weaker values. The network responds as a *MAX* operator, with the maximally stimulated population active proportionally to its input and the rest being quiescent (Eq. 8). **c**. Overview of the possible states of a *P*=2 system receiving intermediate stimulation values 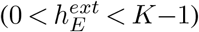. Each tile depicts the intermediate-*ν* solution(s) corresponding to its location on the 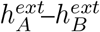 plane (Eqs. 9). Fixed-points (red, green and open circles for 9a, 9b and 9c respectively) are indicated by their positions on the tile (the *V*_*A*_–*V*_*B*_ plane in the boundaries [0, 1]×[0, 1]). The shaded region marks where both stable solutions co-exist, along with the unstable solution. The latter (dis)appears together with the ‘weaker’ stable fixed-point at the boundaries of the shaded region 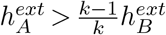 for 9a and 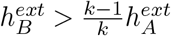 for 9b) in a saddle-node bifurcation. **d**. Initial conditions affect the resultant activity when multiple solutions co-exit. Multiple simulation trials of a *P* =2 network in five different initial states were performed, by varying the fraction of neurons active prior to stimulation onset. Initial conditions are marked on the *V*_*A*_–*V*_*B*_ plane by the serial number of that trial (inset). Also marked is the seperatrix, which divides the plane into basins of attractions of the two stable fixed-points (dashed line). The asterisks (panels c and d) indicate the location of the tile within the zone of co-existence (shaded region in panel c; 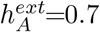 and 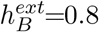).

In both networks and for all *h*^*ext*^ values, the identity of the active neurons keeps changing even as external stimulation is kept constant (panels a and b, rasters). Every neuron thus exhibits the same mean firing rate as every other neuron within each population, a result anticipated by the definition of *ν*_*x*_ as the expected value of *V*_*x*_ for any neuron in population *x* (Eqs. 5, 6). The uniform connectivity implies that the system behaves ergodically, so that the quantity *ν*_*x*_ describes the population-wise mean activity level in *x* as well as individual neurons’ mean rate (panels a and b, bottom graphs).

Neurons in the simple networks above exhibit ongoing fluctuations even though they are all deterministic, receive no external noise, and are connected uniformly within- and between populations. This finding may seem counter-intuitive, but it was reported long ago (Rubin and Sompolinsky, 1989) and is arguably the simplest example of Poisson spiking arising from the balance of excitation and inhibition at the network level. The underlying mechanism is, in essence, the same as in the widely familiar, more intricate version of the model (van Vreeswijk and Sompolinsky, 1996). The inhibitory population tracks the activity of the excitatory population, and returns it as amplified negative feedback. If the inhibitory coupling is strong relative to external inputs (here, if *N*_*I*_*K>N*_EP_[1 +*h*^*ext*^]), it prevents the excitatory neurons from reaching saturation (*ν*=1), forcing each population to settle for only a fraction *ν<*1 of its neurons being active. But there are no pre-determined subsets of active and quiescent neurons: any configuration satisfying the constraints on total activity (Eq. 7, or 8) is valid. There are 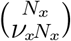 such configurations – an enormous number even for modestly sized networks (eg, the *N*_*E*_=100 neurons in Fig.2(a), have ~10^13^, 10^25^, and 10^29^ valid configurations for *ν*_*E*_ = 0.1, 0.3 and 0.5 respectively). As the network moves between the multitude of configurations, it creates the impression of ‘noisy spiking’ even for deterministic neurons, where the only source of randomness is the update order.

From single neurons’ perspective, a quiescent neuron receives the same total input as those in the active subset: *O*(*N*) excitatory inputs, assiduously counterbalanced by *O*(*N*) inhibitory ones. Neurons will therefore inevitably ‘migrate’ between the quiescent and active subsets as they update their state. Each such migratory event generates an *O*(1) deviation from the balance which is felt throughout the network and, in turn, drives subsequent updates towards (another) configuration satisfying Eq. 7. Activity is thus carried forth in the network by sequences of small perturbations around neurons’ threshold (Rubin and Sompolinsky, 1989; Tsodyks and Sejnowski, 1995), as they remain continually in a high-conductance state (Destexhe et al., 2003). With asynchronous updating, it also follows that neural spiking obeys Poisson statistics.

The persistent fluctuations at zero temperature have a close analogy in physical materials with anti-ferromagnetic interactions (favoring misaligned, rather than aligned magnetic spins). Typically, it is not possible to satisfy all pairwise interactions simultaneously, and the system is then said to exhibit *frustration* (Fisher and Khurana, 1988; Moessner and Ramirez, 2006). This alludes not only to the need to settle for a sub-optimal arrangement of the spins, but also to the lack of a clear, unique compromise to converge on. Instead, there is a profusion of configurations that have a comparable fraction of ‘frustrated’ spin pairs. The dynamical behavior of spins can be described as minimizing an energy (or ‘cost’) function, and the notion of frustration then translates to an energy landscape replete with local minima. Changing from one to another often entails flipping a sizable fraction of the spins, which requires thermal noise (Fisher and Khurana, 1988). But in certain compounds, small clusters of frustrated spins can flip with no cost in energy. These local fluctuations allow the system to wander indefinitely between the multitudes of minimal-energy configurations, even at zero temperature (Moessner and Ramirez, 2006).

The analogy to physical systems of interacting spins has given neural modeling many insights (Hopfield, 1982), but their applicability has been largely confined to networks with symmetric connectivity. Having asymmetric connections – as when excitation and inhibition are effected by distinct neuronal types – precludes a model from being described by an energy function. But PCRNs offer an exception: thanks to the uniform connections to and from the inhibitory pool, a network of N=N_E_ neurons connected by the symmetric matrix **J**^AF^*≡* **J**^EE^ *−K*J^N^ has the same fixed-points as those of Eq. 4 for any *h*^*ext*^*<r*_I|E_*K−* 1 (Rubin and Sompolinsky, 1989). The energy function describing the **J**^AF^ version of the model is equivalent to that of a long-range anti-ferromagnet, conferring a formal basis to the analogy between frustrated spin fluctuations and Poisson firing in the balanced regime.

### Co-existence of Multiple Fixed-Points

Testing all sets of permissible values of Eq. 6 for *P≥*2 reveals that the *MAX* solution (Eq. 8) is not always unique. In a *P*=2 network, the intermediate-*ν* solutions

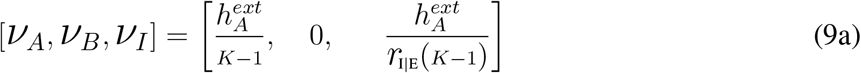

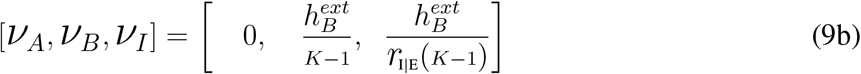

co-exist in a portion of parameter space, and a third fixed-point, given by Eq. 9c, is valid wherever both 9a and 9b exist (Fig. 2c). Stability analysis of the solutions is complicated by Θ being non-differentiable, but can be performed by deriving bounds on *δν*_*x*_ upon local perturbations, or as the limit of large (yet finite) *β* using standard linearization procedures on *G*_*β*_ (Rubin, 1989; Guckenheimer and Holmes, 2013). Both methods indicate that solutions 9a and 9b are stable, whereas 9c is unstable.

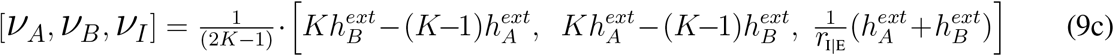

When 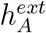 and 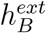 are close in value, a ‘win’ of the more strongly stimulated group is thus still possible, but so is a win of the ‘weaker’ sub-population. Which of the two prevails is determined by the network’s initial state (Fig. 2(d)), with the unstable fixed-point separating the state-space into distinct basins of attraction of the competing stable states (inset in panel d). The weaker attractor’s basin is smaller in size, yet gives it a chance of prevailing when initial conditions are random. This may appear as sub-optimal but, with the addition of spiking noise, it emerges as a central feature of the model.

## 4 Multi-stability at finite temperatures

The intermediate-*ν* states are not an anomaly of the deterministic system. The latter is special for being solvable in closed-form, but numerical solutions of Eqs. 5 for non-zero temperatures reveal continuity with those derived above for Eqs. 6. Each numerically calculated fixed-point starts from one of the closed-form solutions (Eqs. 7, 8 or 9) and gradually departs from it as temperature climbs up, maintaining the same stability (or lack thereof) as its precursor (Supp. Note 2 and Fig. 5), (Rubin, 1989). But while the change to non-zero temperature has little effect on the numeric value of fixed-points, it has a dramatic effect on the system’s dynamical behavior. When multiple solutions coexist, even a slight amount of spiking noise causes intermittent switches between the (stable) fixed-points. With non-overlapping, dense patterns, the switching manifests itself as multi-stable alternations between periods of exclusive activity (‘dominance’) of competing excitatory populations (Fig. 3). In PCRNs with highly distributed and/or overlapping patterns, multi-stability between different attractor states may not be as readily apparent, but it can be discerned with knowledge of the internal representations (Fig.1(b,h)).

**Figure 3:**
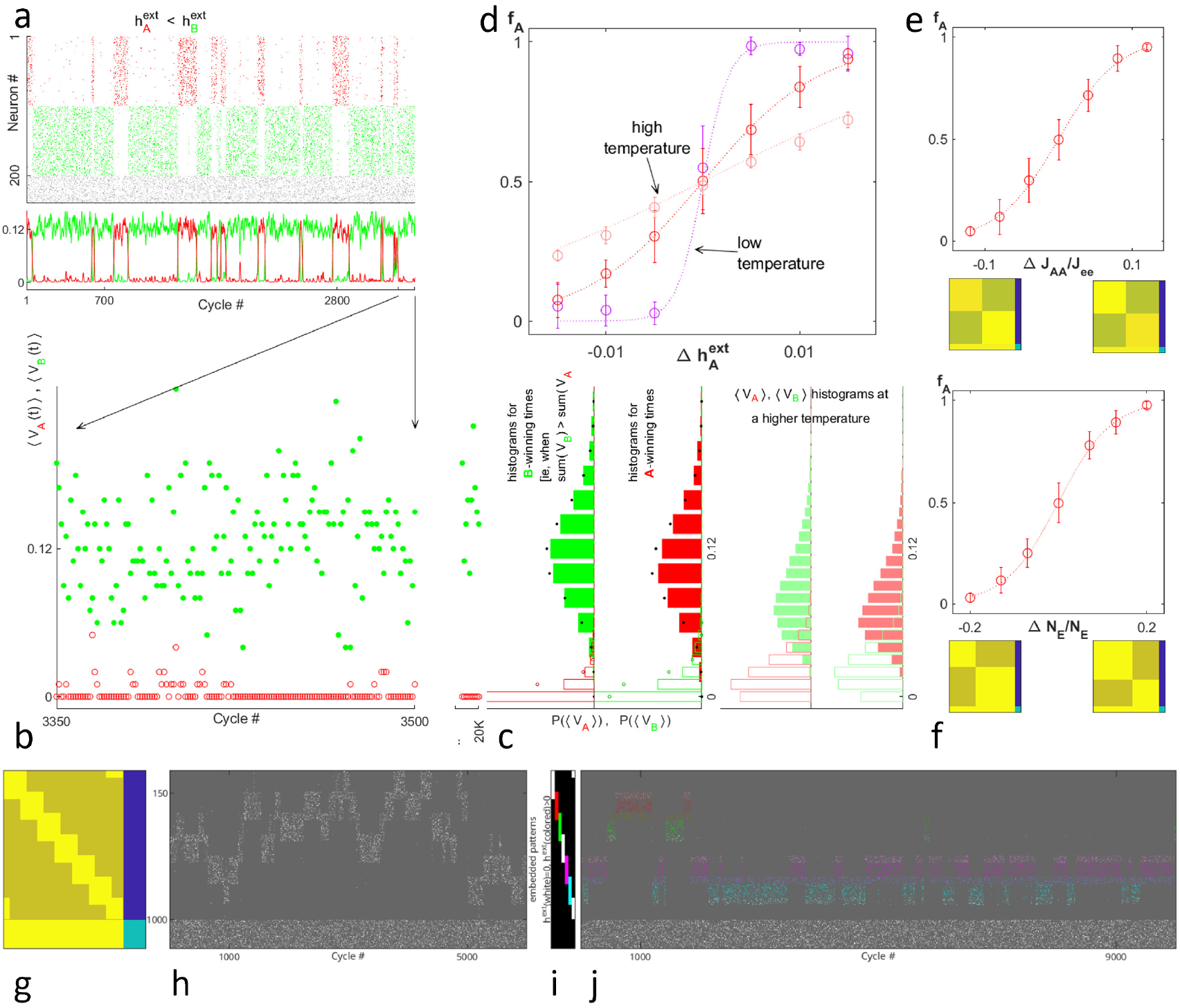
Multi-stability and sampling behavior in PCRNs and in perception. **a-e.** Simulation results in the presence of spiking noise, for a network with two uniformly-connected excitatory sub-populations (*A* and *B*, in red and green respectively) receiving constant external stimulations. **g-j**. Simulation results of a PCRN with seven overlapping sub-populations receiving two kinds of external input. **a**. The competing populations alternate in activity, with the one more strongly stimulated dominating a larger fraction of the time (here, 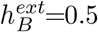 and 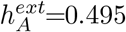). **b**. Zoom-in on the first 300 cycles of the simulation in *a*. Alternations occur when a dip in activity of the dominant population (bold circles) coincides with an uptick in the quiescent population (open circles). **c**. Distributions of mean activity levels for each population, calculated from a simulation run of 10,000 cycles with parameters as in *a-b* (histograms on left), and from a separate run with a higher temperature (right). **d**. The fraction of time *A* is dominant, *f*_*A*_, increases monotonically with 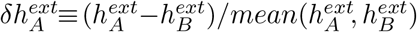. The three curves are for the two temperature values used in *c*, plus a low temperature. **e-f**. Imbalances between *A* and *B*’s fraction-dominance may arise from differences in the internal representations, rather than in external stimulation. Here, the relative strength of internal connectivity within *A* and *B* (panel e) and the size of *A* and *B* (panel f), were systematically varied, while keeping 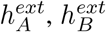 constant at 0.3 (all other parameters as in a-d). **g**. Connectivity matrix of seven embedded block patters of 190 neurons each, with overlaps between adjacent patterns in a ring (ie, blocks 1 and 7 overlap), and an inhibitor pool of 190 neurons. Overlaps occur in about 50% (98) of the neurons in each pattern, 48 neurons in each side, making the total number of neurons *N*_*E*_ = 1, 000. **h**. Time-course of PCRN with neurons in *E* receiving uniform stimulation (*h*^*ext*^ = 0.2), creating alternations between periods of activity of one pattern at a time, rather than neighboring patterns merging their activity, but with alternations moving in waves preferring neighbors. **i-j**. Time-course when external stimulation is limited to two pairs of overlapping patterns (i): pairs 2 and 3 (red and green), and 5 and 6 (magenta and cyan), which don’t overlap. Alternations get trapped within one of the pairs for a while before transitioning to a pattern in the other pair and getting trapped there for another while (j). A similar structure of inhomogeneous transition probabilities has been reported in perceptual experiments with a quad-stable binocular rivalry stimulus (Suzuki and Grabowecky, 2002), where observers had long periods of ‘trapping’ within each pair of related percepts, and alternated between pairs far more infrequently.

Each competing population enjoys some periods of dominance, but more strongly stimulated ones prevail a larger proportion of the time (Figs. 1(e), and 3(a)). Mechanistically, this comes about via an asymmetry in the probability of switching. Due to spiking noise, the total number of active neurons in each population is no longer constant, but rather hovers about its mean with a binomial distribution (Fig. 3(b)). Alternations happen when the dominant population dips markedly below its mean, just as a quiescent population surges well above its own. In a *P*=2 PCRN, the probability this occurs is given by the overlap of the histograms of activity levels in the dominant and quiescent populations (panel c, filled and open bars respectively). The weakly stimulated sub-population is less active, on average, during dominance periods (filled bars, red vs green) as well as during quiescence (open bars), resulting in greater overlap when it is dominant (filled red bars) and thus higher probability of switching back to its stronger competitor. The fraction of time *A* prevails, denoted *f*_*A*_, increases monotonically with the relative strength of its stimulation, in a sigmoidal manner that echoes individual neurons’ probabilistic response function (panel d). The shallower slope of 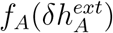 at higher temperatures, mediated by reduced asymmetry between the probability of switching to-and from *A* as its overlap with *B* grows (panel *c*, histograms on right vs left), further underscores the link between spiking noise in individual neurons and behavior at the system level.

Multi-stability thus mediates probabilistic sampling: where the deterministic system gets stuck in one response regardless of comparable alternatives (or even better ones; Fig. 2(d), trial 5), the PCRN ‘hedges its bets’ by alternating stochastically between competing responses, spending larger portions of the time in those more strongly supported by the input. A similar behavior is found for perceptual bi-stability of ambiguous stimuli, where parametric manipulations that strengthen one possible interpretation over the other produce a gradual, monotonic increase in the probability of that percept to dominate (Levelt, 1965; Blake, 2001; Hupé and Rubin, 2003). Moreover, the dynamics of alternations are consistent with a Markov process with stationary statistics: the mean and variance of dominance durations for each competing percept are stable over time, with no systematic correlations between percept durations in subsequent epochs (Fox and Herrmann, 1967; Rubin and Hupé, 2005). And yet, viewing the alternations as arising from noise-driven sampling has so far, faced a major obstacle, in turn reopening the challenge to explain the non-monotonic distributions of percept durations (Moreno-Bote et al., 2007, 2008, 2010).

Rather than contemplating a process that is fundamentally stochastic in nature, the bulk of modeling work on perceptual bi-stability has therefore posited alternations are caused by a process of ‘fatigue’ in the winning population: spiking adaptation, synaptic depression, or both (Lehky, 1988; Laing and Chow, 2002). In that scenario, the peak in the distribution of percept durations results from a periodic process – oscillations between the two percepts – due to the predictable time-course of the fatigue process, with auxiliary noise sources adding variability to broaden the peak. But there are problems with this idea, too. The long time constants of the fatigue processes predict systematic effects on the statistics of the durations, in particular non-stationary means and second-order correlations, which auxiliary noise cannot fully mask. Yet such effects are not observed experimentally (Rubin and Hupé, 2005). While commonly demonstrated with visual stimuli (Levelt, 1965; Blake, 2001; Rubin and Hupé, 2005), bi-stable perception occurs also in other sensory modalities (auditory, tactile, olfactory (Schwartz et al., 2012)) and thus seems emblematic of how the brain responds to deeply ambiguous stimuli.

Although previous network models based on mutual inhibition have been shown to capture many characteristics of the dynamics of perceptual alternations (Laing and Chow, 2002; Lehky, 1988; Moreno-Bote et al., 2007), a few unresolved issues have persisted. At first glance the bi-stability in Fig.3(a) may seem like alternations between two competing network states, like those familiar from many models of perceptual bi-stability and decision-making. But there is an important difference: previous models have utilized traditional ‘winner takes all’ behavior, with neurons in the winning population firing at or very near their maximal rate (Laing and Chow, 2002; Wang, 2002). In contrast, here the winning neurons fire at only a fraction of the maximal rate. Thus, a ‘win’ of one population, say *A*, no longer corresponds to just one network state (‘all *A* neurons active’). Rather the number of possible configurations the system may be has as lower bound, 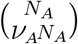, the number of configurations at zero temperature. Hence, while at the population (‘macro’) level an alternation seems like a single transition between two competitors, in reality it depends on the coincidence of many single-neuron (‘micro’) events. The dominance durations correspond to the intervals between those rare coincidences and, for large N, their distributions indeed resemble a skewed Gaussian, aligning with experimental observations (Blake, 2001; Rubin and Hupé, 2005).

The dependence of *f*_*A*_ on *δh*^*ext*^ comes about via the latter’s effect on the relative size of *A*’s basin of attraction. While this may seem but a nuance in the simple case considered here, it is relevant for understanding probabilistic sampling in more complex cases of competition between multiple, overlapping attractor states and/or patterned stimulations. In such cases there may no longer be a simple scalar quantity like *δh*^*ext*^ to derive, but the concept of a basin of attraction for each of the competing network states still holds. *f*_*A*_ reflects the relative strength of the *internal representation* of percept *A* (rather than that of the external, physical, stimulus), that is, the size of its basin of attraction (relative to those of the competing attractors). This can arise from the strength of the internal connectivity in A (Fig.3(e)), the size of the population (Fig.3(f), and overlaps between A and other percepts (Figs.1(f-h)). The hypothesis that *f*_*A*_ depends on the relative size of *A*’s basin of attraction in those cases, too, gains support from findings in perceptual bi-stability. A recent re-analysis of a large corpus of published data has uncovered a systematic relationship between *f*_*A*_ and the mean dominance durations of the two percepts (note those are measured in units of time, not as fractions; (Rubin and Talbot, 2016)).

Given a stimulus configuration for which *f*_*A*_ =0.5, denote its mean dominance durations by 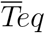 (for ‘equi-dominance’) and define 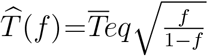. For all other configurations in the same experiment (ie, those which yielded *f*_*A*_ values other than 0.5), the mean durations of percepts *A* and *B* closely followed the functions 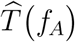 and 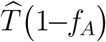, respectively. This hitherto overlooked dependence of the mean durations of both percepts on *f*_*A*_ (or, equivalently, on *f*_*B*_ =1 *f−*_*A*_) was found to hold for numerous binocular rivalry experiments which manipulated the stimulation strength to each eye separately (Levelt, 1965; Blake, 2001), as well as for bi-stable stimuli in other perceptual domains, where it is not experimentally feasible to affect the strength of just one of the two competing percepts because they share the same sensory stimulation (eg, ambiguous motion displays; (Moreno-Bote et al., 2008, 2010)). And all these variations are achieved without the need of a non-markovian model.

Furthermore, multi-stable behaviors, like those shown in Fig. 1(d,e) of a *P* = 4 block diagonal PCRN can now be understood simply by extending the foregoing analysis to *P >* 2. Moreover, simple modifications to purely block-diagonal connectivity can guide the understanding of PCRNs with more intricate **J**^EE^ matrices, including those embedding continuous patterns. When a set of block patterns are modified to add overlap between adjacent pairs (Fig. 3(g)), the connectivity matrix broadens from block to multi-diagonal. With flat stimulation, the overlap terms bias alternations towards each pattern’s nearest neighbors, resulting in a ‘wave’ global activity (Fig. 3(h)). When two pairs of adjacent blocks that have no direct overlap are stimulated (Fig. 3(i)), alternations become ‘trapped’ in one pair for prolonged periods before switching to the other pair (Fig. 3(j)). Rich changes in the (formerly flat) Markov transition matrix can thus be obtained by changed stimulation as well as embedded patterns. The Gaussian connectivity (Fig. 1(f)) can be thought of as the limiting case of a smoother, multidimensional matrices, giving rise to wavy patterns similar to those observed during spontaneous activity in V1 orientation columns (Supp. Fig. 15 from (Kenet et al., 2003)). Conversely, in response to range-limited boxcar stimuli the same network performs ‘peak selection’ while also recovering the shape of the original embedded pattern, as observed in physiological data (Supp. Fig. 14 from (Wu et al., 2008)). A permutation exists in the network matrix of random sparse patterns (Fig. 1), where subgroups of neurons can be found each of which possibly overlapping some other subgroups and yet the system doesn’t get confused (ie, merge patterns). With flat input, neural activity is sustained in one pattern before switching. Similar to Hopfield networks, a PCRN non-linearly selects patterns, adhering to a calculable storage capacity. But, unlike them, it indicates multiple pattern matches by alternating selections at a frequency related to match likelihood, instead of maintaining the strongest signal. The response range of a PCRN is determined by the external stimulation while the tuning is shaped mainly through intra-cortical connections, as is the separation ability. Moreover, as temperature is turned on and increased, simulation results confirm the symmetrized system behaves qualitatively similar to the PCRN (Supp. Fig. 11), solutions in the two systems may depart numerically, but not substantively. This, in turn, suggests that the system may be sampling competing states according to the Boltzmann distribution (Hinton and Sejnowski, 1983), not just for simple, block-diagonal matrices but also for those embedding patterned and overlapping attractor states, and it will be of further value for explaining the system’s sampling behavior at non-zero temperature.

The ability of the PCRN connectivity formulation to recover the shape of continuous variable patterns relies on the binary neurons’ activities being well below saturation, so as to assume varying values of *ν*_*x*_, which in turn relies critically on the networks operating in the balanced regime. Due also to balance plus frustration, each set of neurons belonging to the same attractor are grouped into a binary unite that exercises the function of an analogical-probabilistic unit when in the balanced regime at finite temperatures, opening a path to PCRN computing machines.

## 5 Compound PCRNs

The focus so far has been on ‘deep’ ambiguity scenarios requiring choice among limited alternatives. However, multiple fixed-points coexistence cones are rather narrow (Fig. 2(c)), explaining the rarity of bi-stability which requires uncommon environmental conditions. Despite numerous sensory interpretations, deep ambiguities are infrequent, suggesting that competition is often resolved by a single attractor without sampling. Nevertheless, the brain samples local ambiguous cues in milliseconds, forming conscious perceptions over seconds. Various probabilistic cue combinations yield several compound probabilities (Hoyer and Hyvärinen, 2002), which inform further modules to seek a globally consistent interpretation of the signals. Calculation of the ‘majority vote’ of a set of probabilistic cues is the simplest form of cue combination. The feed-forward PCRN architecture in Fig. 4(a) performs this computation for a binary decision; generalization to *P >* 2 is straightforward. Independent PCRNs provide cues to decide between two options (colored red or green). Each PCRN delivers input through excitatory connections from its red and green subpopulations to the corresponding subpopulation of a single output PCRN. All input PCRNs receive uniform, equal external input, but a preference in favor of red is created through **J**^EE^ by increasing the interconnection strength among neurons in red subpopulations and decreasing it in green subpopulations. The **J**^EE^ of the output PCRN is kept unbiased. Simulations show that *f*_*R*_, the frequency of time red prevails in the output PCRN, is always larger than the probability summation alone of finding red winning in the majority of the input PCRNs. The effect is caused because when the red subpopulation in any input PCRN dominates, its mean activity, *V*_*R*_, is slightly larger than when its green subpopulation dominates, *V*_*R*_ *> ν*_*G*_ (see Supp. Fig. 7). This in turn makes the input to the output PCRN’s red subpopulation larger than the input to the green subpopulation any time the number of red winners is larger than or equal to the number of green winners in the input layer. Note that the difference between mean activities is essential, particularly braking ties. This is an asymmetry that can’t be captured by single binary units such as spins of the Ising model. The appeal of a network that can do probabilistic sampling and make probabilistic calculations has been suggested long ago (Hinton, 2007; Griffiths et al., 2010; Pearl, 2014). Yet the idea has never been realized as it’s done here within the constraints of a biologically plausible neural model. Moreover, the approach is supported by and aligns with the evidence that cortical sensory variability is not a reflection of noise but of uncertainty in the stimulus (Hoyer and Hyvärinen, 2002).

**Figure 4:**
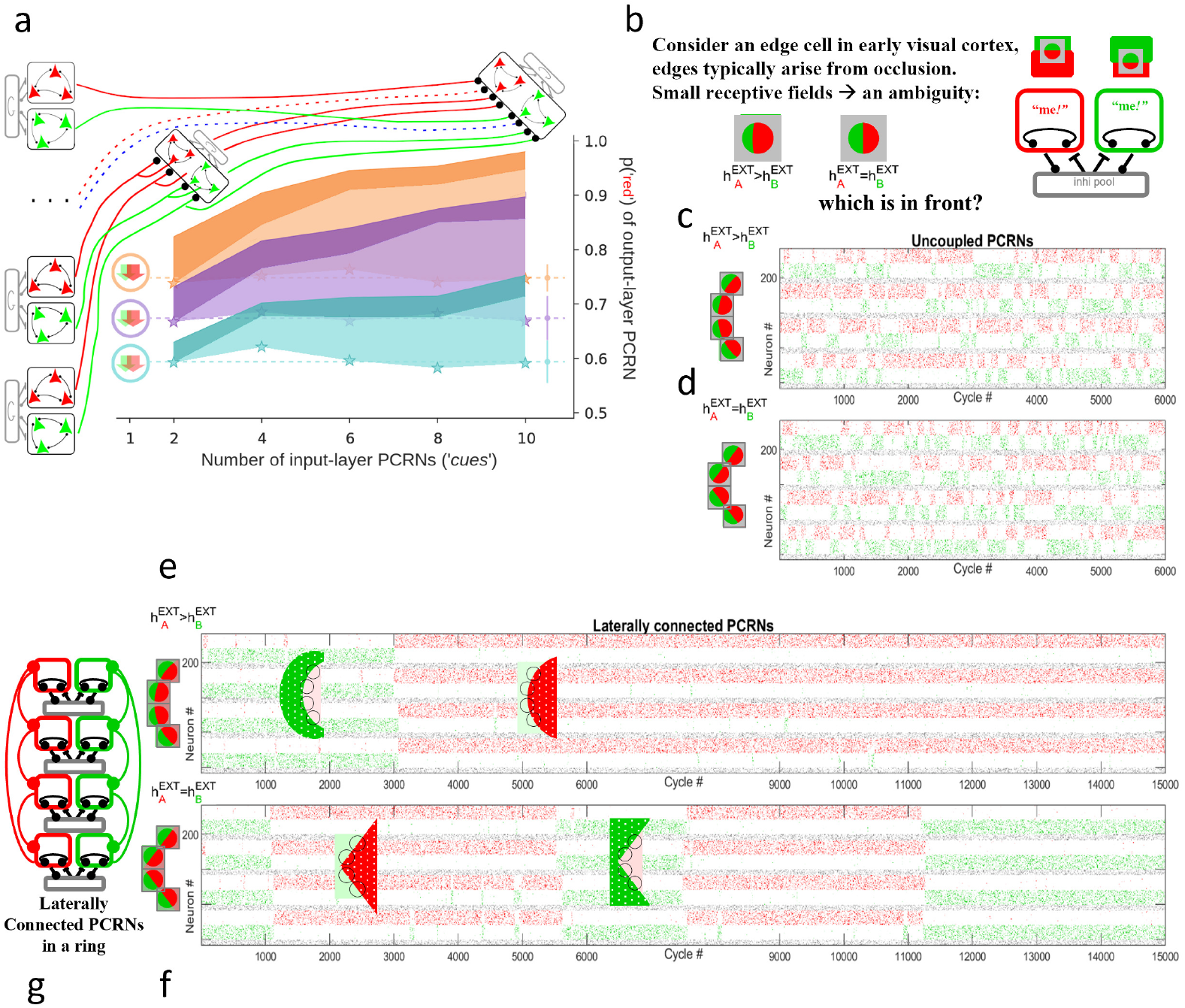
Feed-forward and laterally connected compound PCRNs. **a.** Probabilities computed from 50,000 cycle simulations over three variants of five feed-forward PCRN architectures, each combining an even number of cues - from 2 to 10. Schematic models of 2- and 10-cue architectures are shown. The input layer on the left are identical *P* = 2 PCRNs with the interconnection strength in the *A* (red) subpopulation slightly higher than in the *B* (green). Three strengths yielding probabilities, *p*(*A*), of cue *A* winning of approximately 0.6, 0.67 and 0.75 (colored blue, purple and orange respectively) are the variants. The pale area is the probability of finding *A* winning in the majority of the input PCRNs. The vivid color is the probability of finding *A* winning in the output layer PCRN, always above the pale area. External stimulation is flat. The stars indicate the probability of finding *A* winning in an input layer PCRN. Above the plot are the 2-cue (left) and the 10-cue (right) output PCRNs. **b-g**. Simulations of two pairs of arrays of four PCRNs, one without and the other with lateral connections, receiving input under two different conditions from contiguous visual receptive fields. The fields are presented with two surfaces that create an edge. In one case, the edge is curved (panel b, most left), in the other, the edge is straight (panel b, next cell to the right). Each PCRN must decide which side is figure, and which is ground (panel b, PCRN on the right). **c,d**. Two time-courses of unconnected PCRNs, each a copy of the PCRN depicted in panel (b), receiving input from receptive fields according to the figures to the left of the time-courses. **c**. Because 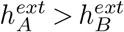, *A* tends to dominate in all four PCRNs (ie, red is in the foreground). However, finding all four agreeing on the same foreground is rare. The edges in their fields are created by a circle. **d**. With no dominant global subpopulation, there is no globally consistent state where all PCRNs agree on figure and ground. The edges in the fields are straight lines formed by the rectangle’s corners, hence, 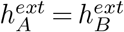. **e-g**. Modeling figure-ground perceptual grouping: time-courses of 4 PCRNs with inputs as in panels (c) and (d), but laterally connected as in panel (g). Lateral connections drastically change the dynamics: PCRNs quickly synchronize, and both simulations (by chance) first reach a globally consistent state where the green surface lies above the red. They then randomly alternate between globally consistent states while maintaining synchronized PCRN state transitions. These global states become attractors of the compound system, and transitions require coincident similar local events across the 4 PCRNs, leading to long durations—longer in panel (e) because, as in panel (c), 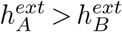, so *A* tends to dominate. Further details on laterally connected PCRN dynamics appear in Supp. Figs. 8-9.

**Figure 5:**
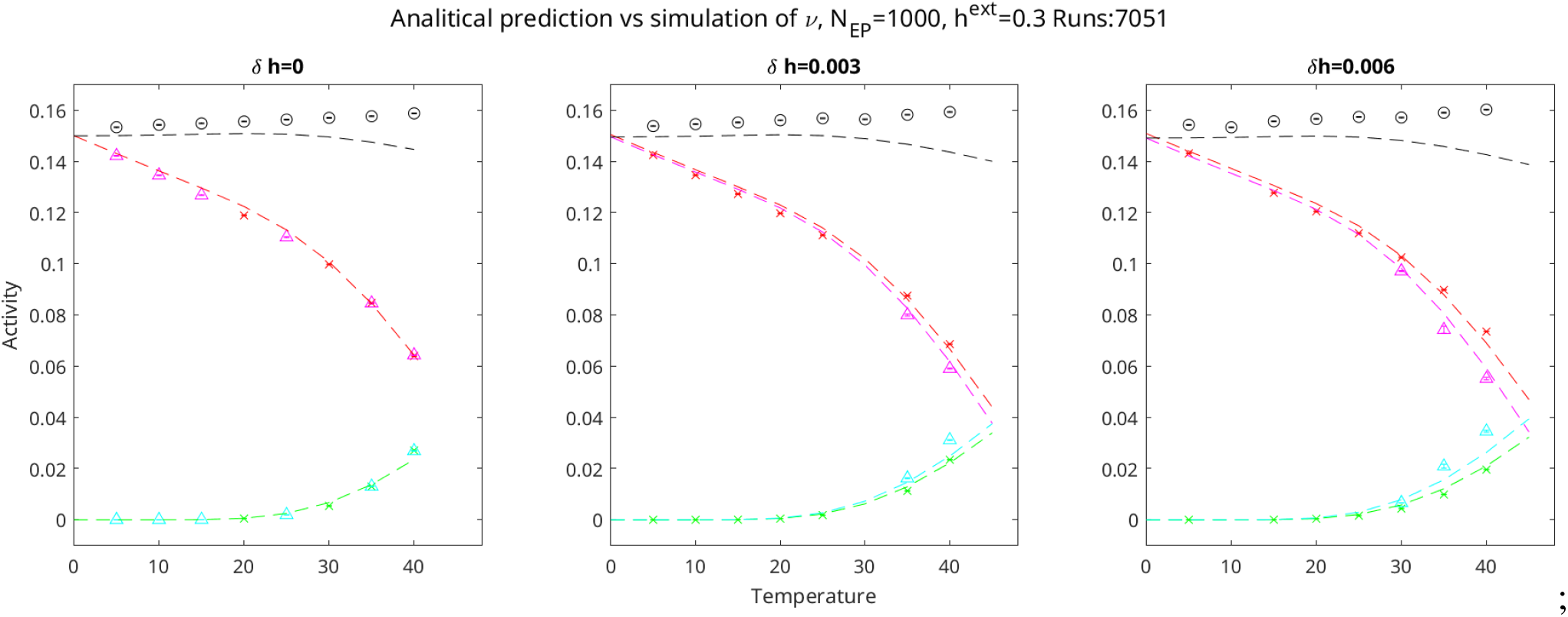
Experimental evaluation of analytical approximation of *δν*. Simulation results of a *P*=2 PCRN with 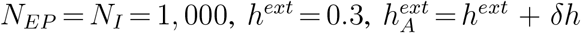, and 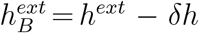. The size of the subpopulations was increased for accuracy. Temperatures were adjusted accordingly. Crosses mark the mean activities when A wins: *V*_*A*_ is a red cross, and a green cross is the activity of B when losing which rises from 0 as the temperature increases. Triangles mark the activity when B is winning: *V*_*B*_ is cyan, and magenta the activity of A. The dashed lines are the predicted activities, red or magenta when A or B is winning, cyan or green when A or B is losing. For *δh* = 0, the lines overlap, but they separate as the temperature increases when *δh* = 0, and *V*_*A*_ *>ν*_*B*_. There are no alternations when the temperature is below 30. The black line and circles correspond to the activity of the inhibitory pool.

**Figure 6:**
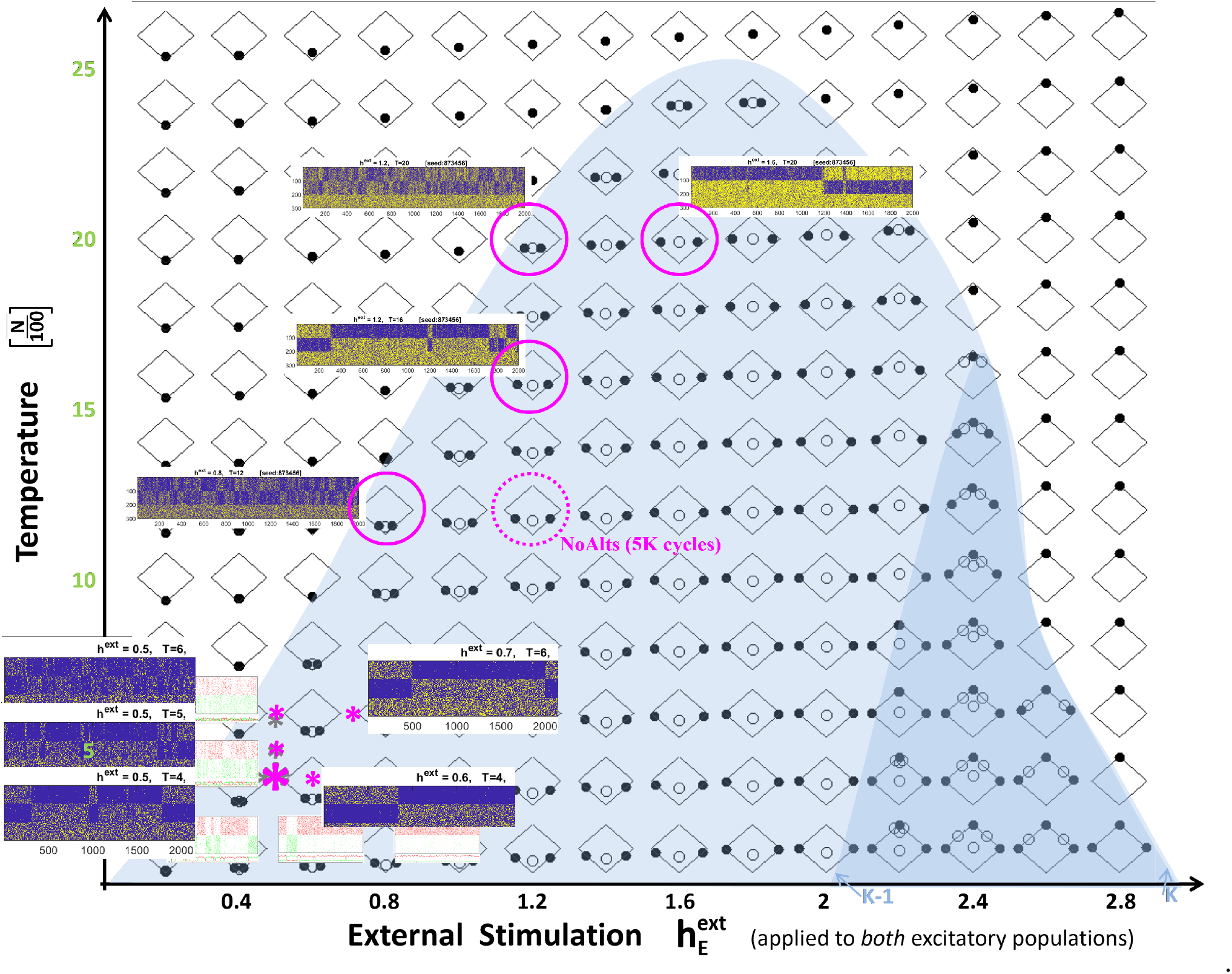
Balanced inhibition unleashes sampling by curbing mean firing rates. Phase diagram of a PCRN embedding *P*=2 binary patterns, with the region where two stable solutions coexist demarcated on the stimulation-temperature parametric plane in light-blue shading. Fixed-points of Eqs. (6) were computed numerically for each combination of stimulation 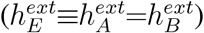 and *β*, and marked on the tile in the corresponding location. (Conventions as in Fig. 2b, but here tiles are presented as diamonds to accentuate the solutions’ symmetry.) As 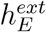 grows, the stable fixed-points increasingly resemble traditional winner-take-all, with [*V*_*A*_, *V*_*B*_] approaching [1, 0] or [0, 1] for all but the highest temperatures. (Note the close agreement between the numerical solutions at low temperatures and those derived in closed-form for zero temperature, where saturation is reached at *K−*1; cf bottom row here with main diagonal in Fig. 2(c)). Simulations showing a marked slow-down of alternations as external input is increased from 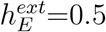 to 0.6 in the two raster in the bottom row, with temperature held at 0.04, and an increase from 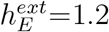 to 1.6 in the two raster at the top, with temperature held at 0.2. Network simulations illustrating the effect of increasing temperature on alternations between the competing populations is also shown. External stimulation is held at 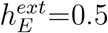 while temperature is raised from 0.04 to 0.06 (left column rasters). The steep parametric effects (ie, the ‘crowdedness’ of the asterisks) are due to the small *N*; as *N* grows, the effects become more well-spaced but simulations will take much longer.

**Figure 7:**
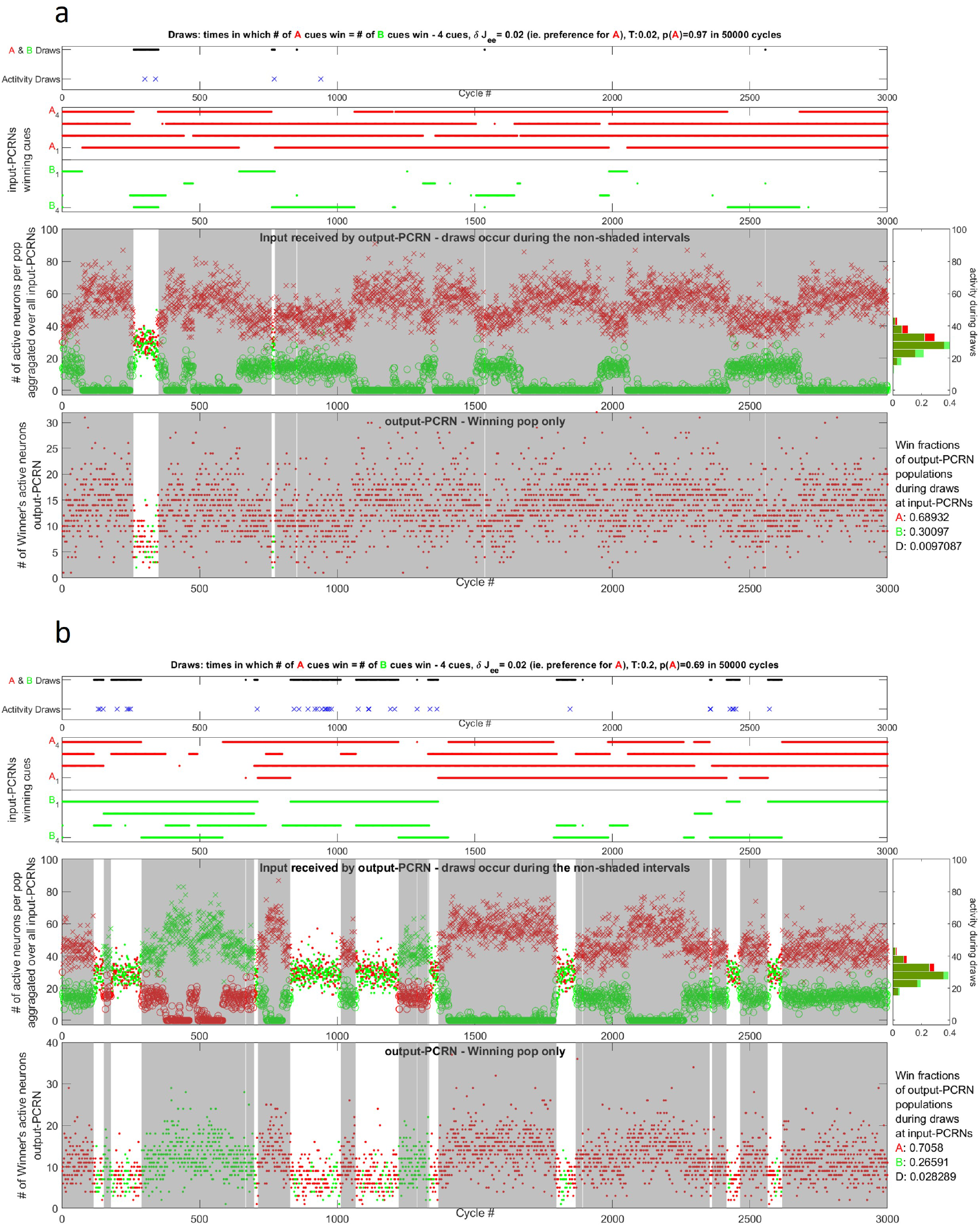
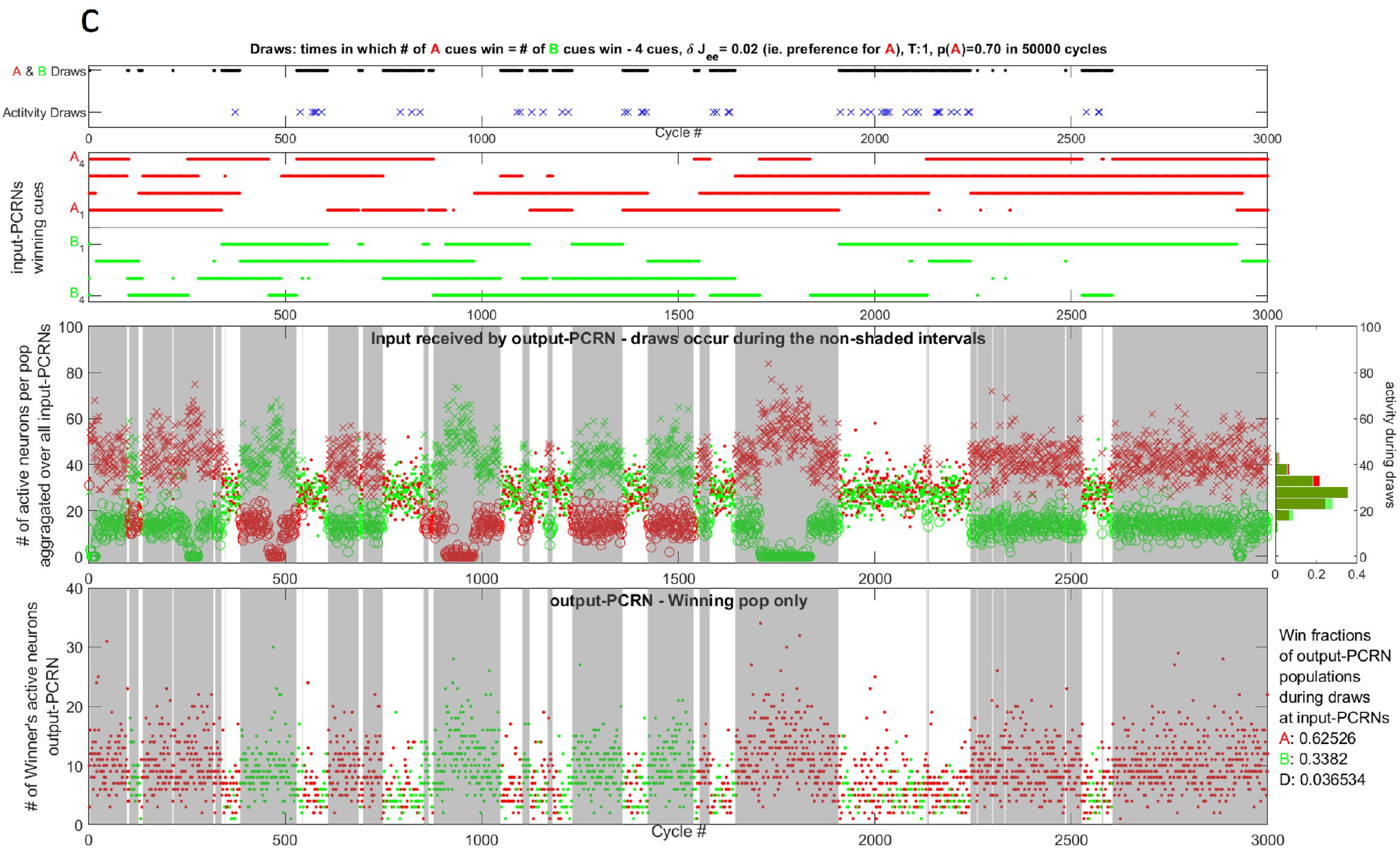
Effects of ties in outcomes of a 4-cue feed-forward network. **a-c.** Results, spread over two pages, of simulations running a 4-cue biased network at three different temperatures, 0.02 (panel a), 0.2 (panel b) and 1 (panel c), and a single *δ***J**^EE^ = 0.02, that yields *p*(*A*) = 0.6. All the panels show time-courses of the first 3,000 cycles out of 50,000 cycle simulations. The top time-course in each temperature group shows the points in the simulation when there is draw (black line) and of those, the ones for which the total number of active neurons in the A subpopulations is the same as in the B subpopulation (blue crosses). The next time-course shows the winning population for each of the four input PCRNs. The last two time-courses show the number of active neurons in the input PCRNs and in the output PCRN respectively. The shaded areas are over the intervals where there is majority of winners in the input PCRNs for one of the two populations. It is clear that there is a larger chance to get a draw as the temperature increases. But the activity in the input PCRNs seems to show slight bias towards higher activity in A than in B, implying that the input to the output PCRN is higher in the A subpopulation. This is corroborated by the histogram of activity on the right. Higher values are shown by A (red) than B (green). To the right of the output PCRN time-course are listed the probabilities of finding the PCRN in a state where A is winning, or B is winning or in a draw. By the nature of the XOR enforcement of the PCRN, the draws are kept below 4% in spite of the high fluctuations in its input. But, more importantly, *p*(*A*) for the full simulation of 50,000 cycles is 0.97 for temperature 0.02, 0.69 for temperature 0.2 and 0.70 for temperature 1.

Feed-forward compositions set aside, PCRNs truly excel when addressing ‘weak’ ambiguities. These arise from fragmented information provided as input, with each fragment being ambiguous on its own, yet collectively resolved into one or two consistent interpretations. Fig. 4(b) shows an illustrative example in which ambiguous input arises from receptive fields that detect the edge created by two surfaces at different depths. A PCRN can’t locally decide between the red over green surface or vice-versa. PCRNs receiving adjacent edge continuation inputs will switch between interpretations independently, despite the inconsistency of assigning the same surface to different depths simultaneously. To achieve global consistency, adjacent PCRNs cue each other for state synchronization. While feed-forward PCRNs can combine cues, this scenario favors reciprocal connectivity, with a single layer of PCRNs having lateral excitatory links between same-feature sub-populations (Fig. 4(g)). Lateral connectivity exhibits more dramatic effects than feed-forward: it effectively synchronizes and slows alternations, and enhances activity (Supp. Fig. 8). Figs. 4(c-f) show examples: a red semi-circle in one example and a rectangle corner in the other on a green surface. Subpopulation *A* receives stronger stimulation in the semi-circle scenario, and both are equally stimulated in the rectangle. Figs. 4(c,d) present baseline rasters of four uncoupled PCRNs receiving this input. There are 16 global states, but a semi-circle bias increases the likelihood of PCRNs of Fig. 4(c) to be in global states with more *A* than *B* winners; globally consistent states are rare. One occurs around cycle 1100, with *A* dominating. Fig. 4(d) lacks visible consistency. In contrast, laterally connected PCRNs of Figs. 4(e,f) quickly achieve and alternate between consistent states, with transitions through inconsistent states happening too quick to detect. Both have long durations of consistency, but in Fig. 4(e), external inputs cause longer ‘*A* wins everywhere’ states than ‘*B* wins everywhere’. The rapid transitions between globally consistent states are linked to, yet distinct from, why *f*_*R*_ in the output PCRN of the feed-forward network exceeds predicted probabilities: the behavior results from differences in mean activities rather than static biases in network parameters. The external input to a PCRN depends on adjacent winners. If both neighbors have the same winner, say *A*, subpopulation *A* receives input from both the receptive field and the mean activity of the *A* subpopulations of its neighbors (modulated by the strength of the lateral connections). Subpopulation *B* only receives input from the receptive field. This variation, *h*_*A*_ *> h*_*B*_, results in dynamics where *ν*_*A*_ *> ν*_*B*_, steering the PCRN towards *A* wins (see Figs. 3(a-c)), aligns with neighbors, and further increasing *ν*_*A*_ for all three. This inter-dependency favors globally consistent states (Supp. Fig.8). As with the feed-forward PCRN, differences in subpopulation activities reflect frustration between neighbors at different strengths not experienced by a ‘1’ spin next to a ‘-1’ spin in an Ising model.

**Figure 8:**
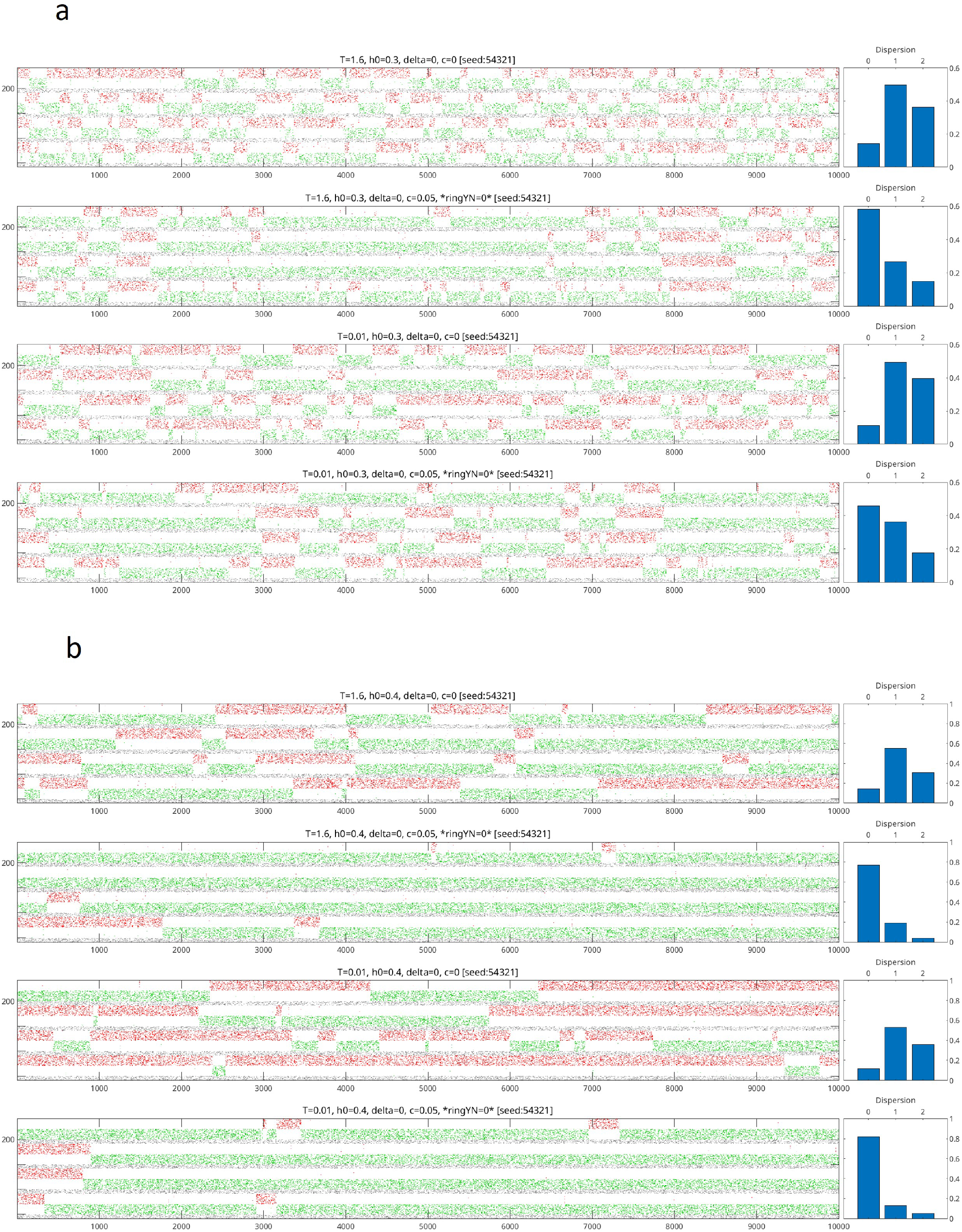
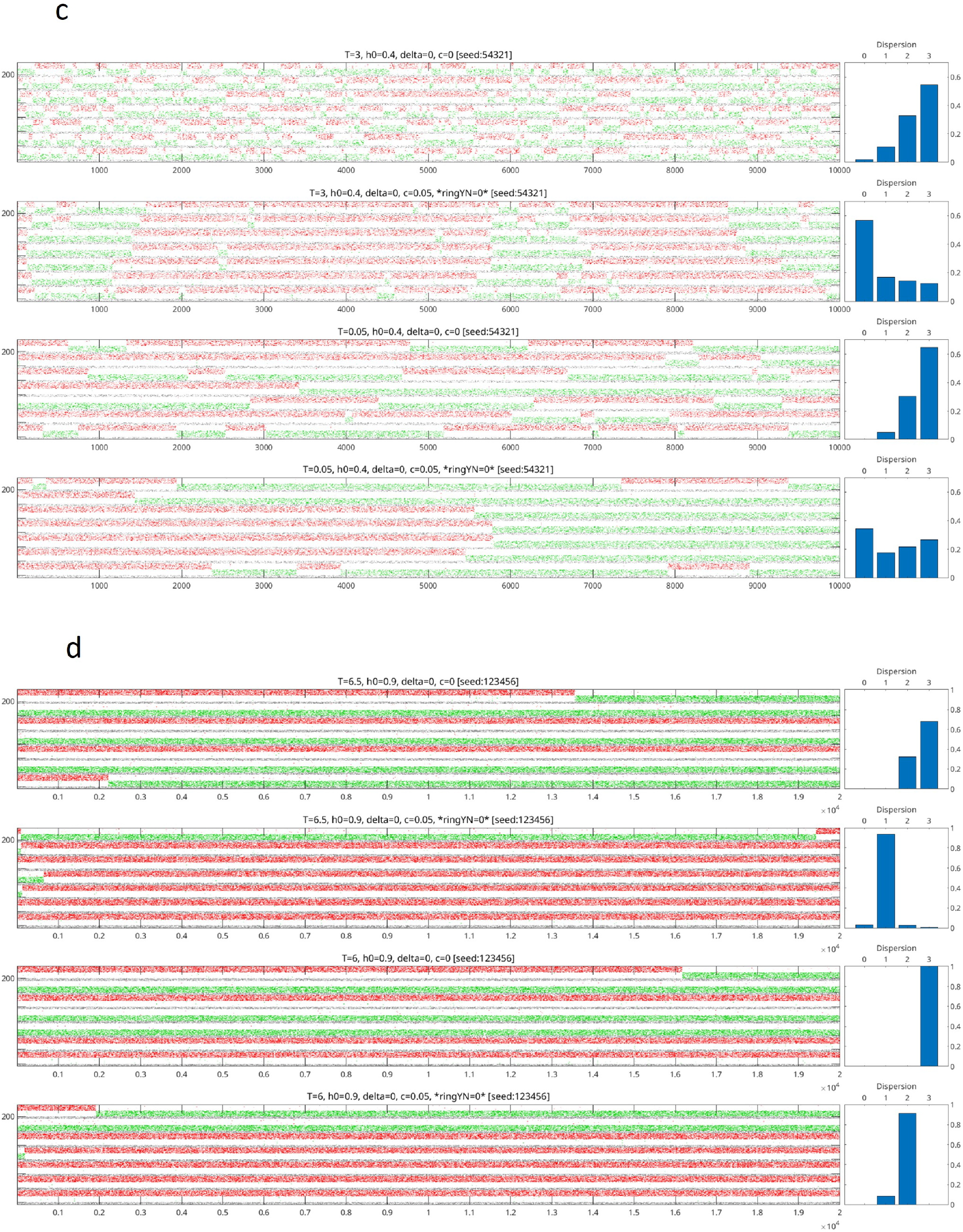
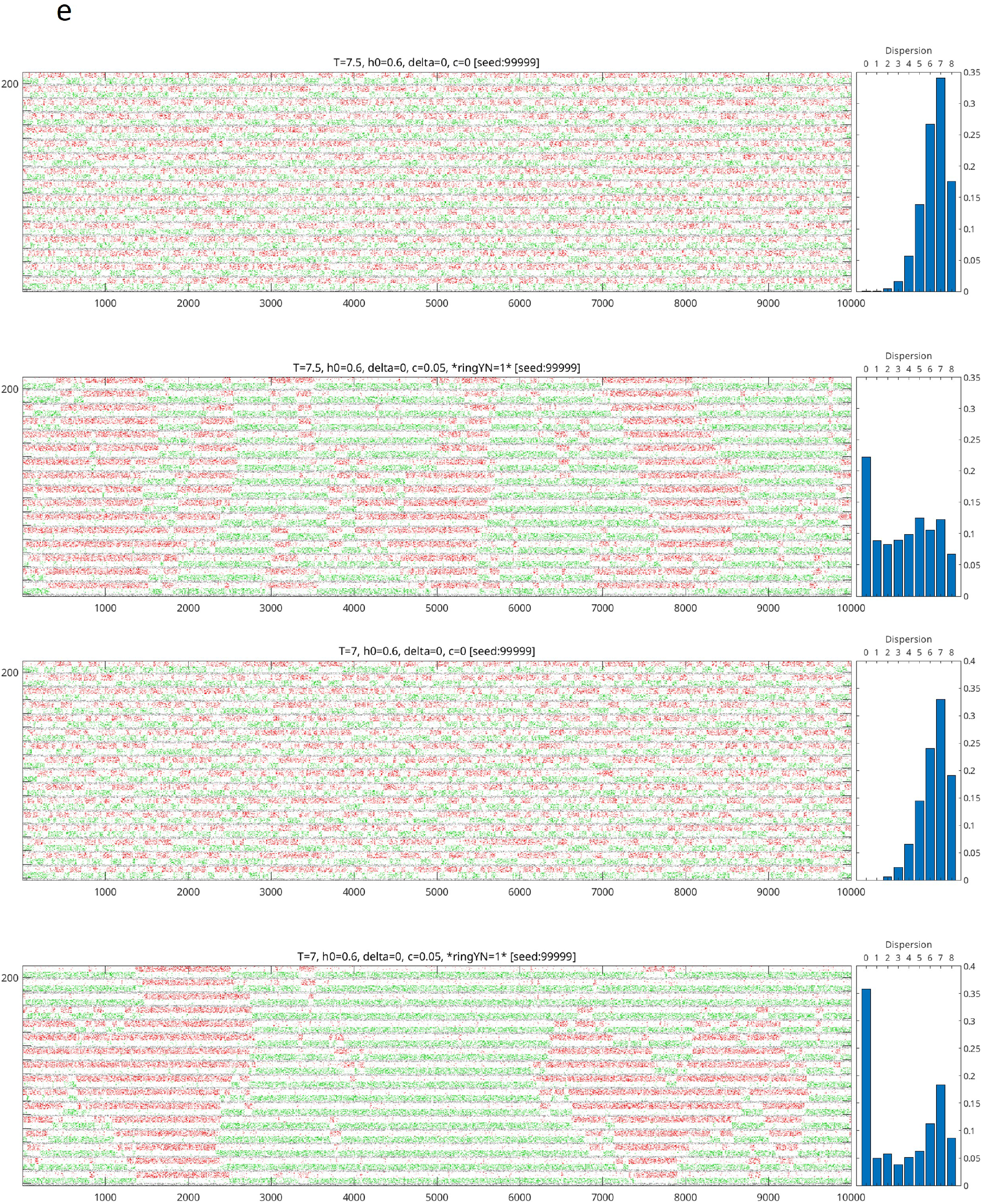
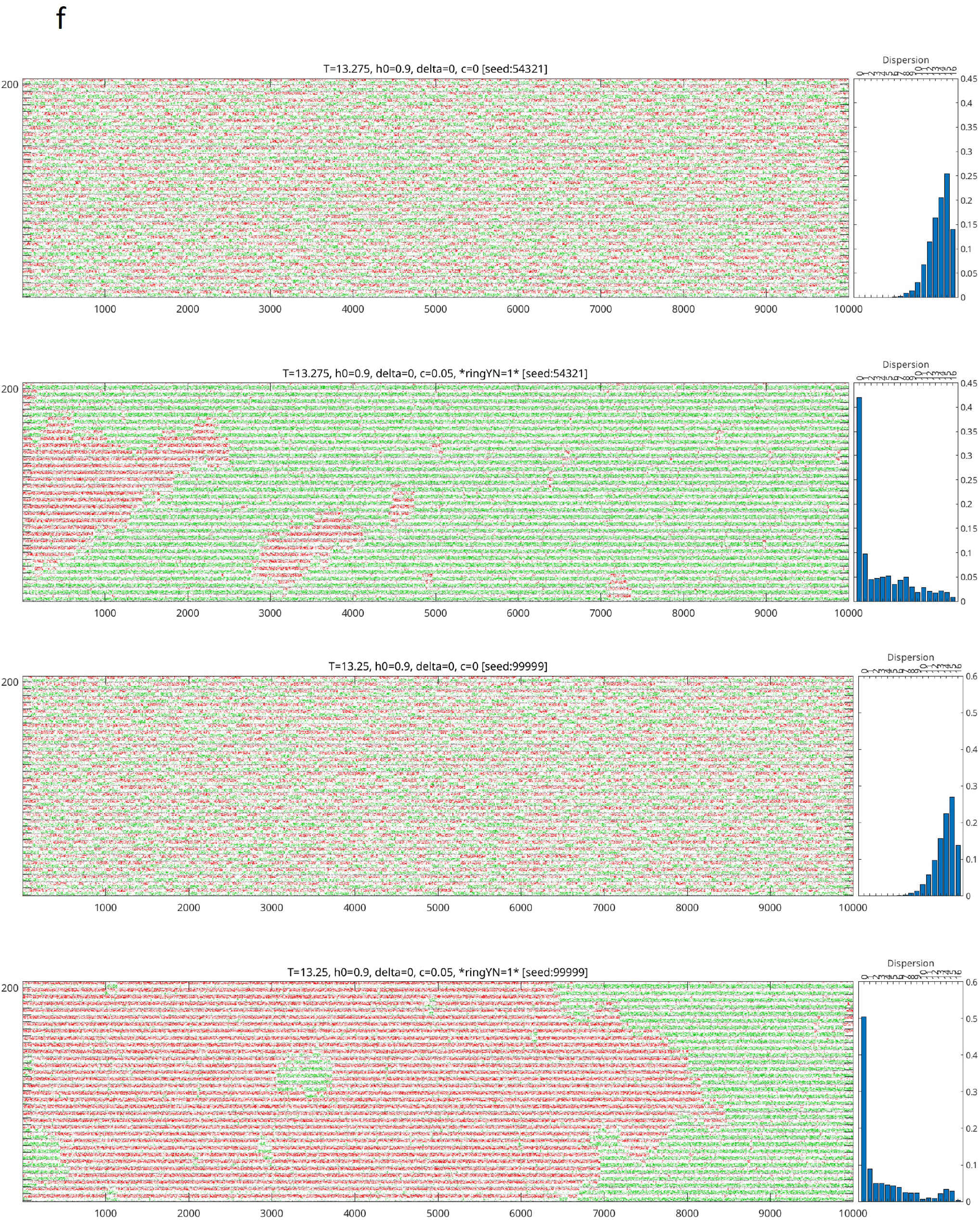
Effects on variability of basic PCRN lateral connections. **a-f.** Spread over four pages, unconnected and laterally connected (strength *c*) PCRNs with no biases (*δ* = 0) in *h*^ext^. Simulations used the same code as in Figs. 4(d,f), but with varying temperatures, external inputs, and sizes. **a**. Four time-courses of sets of four *P* = 2 PCNRs, unconnected in time-courses 1 and 3, and laterally connected in 2 and 4. All get the same external input, *h*^*ext*^ = 0.3. The temperature, *T*, is 1.6 in the top two, and 0.01 in the bottom two. Lowering *T* slows down the transitions in the unconnected PCRNs. This makes more difficult the convergence to a global stable state without local transitions. In contrast, this convergence is observed in the connected PCRNs, with larger durations at the high temperature. The connected system tends to converge to global states where the dominant subpopulation is the same in most of the individual PCRNs. Next to each time-course, there is a bar-chart that groups the states by dispersion: the minimum number of local PCRN transitions needed to get a global state where the same subpopulation dominates in all the PCRNs. The probability of getting to such a global state is 1*/*8, if there are no biases. At *T* = 1.6, the chance is close to 60% in the compound PCRN, and close to 50% at *T* = 0.01. That probability is 0.11 for the unconnected PCRNs at the low temperature and 0.1413 at the high temperature. **b**. Increasing only, the external input, *h*^*ext*^ == 0.4, has the same overall effect of slowing down transitions. The qualitative properties between the different cases stays the same, but the chances of getting to a 0 dispersion global state is now 0.81 for the low temperature and 0.77 for the the high temperature. **c-d**. Increasing the number of PCRNs needs higher external inputs (panel c). And it also requires higher temperatures. The input must be such that alternations should occur in the unconnected PCRNs; otherwise, a significant increase in *h*^*ext*^ a globally consistent state of 0 dispersion might required much more time to reach (panel d). **e-f**. Two simulations of a 16 and two a 32 compound PCRNs are shown. Both PCRNs require higher external inputs and temperatures to reach globally consistent states. They show how alternations moving in waves, to the point, that one can’t see separate patches of the same color appear and progress to a state where the patches grow and merge.

Compositionality can be further demonstrated through the modeling of orthogonal-grating monocular rivalry, which employs a 2D network of laterally connected PCRNs (Supp. Fig. 9). PCRNs are presented with red and green stimuli from receptive fields arranged in overlapping red columns and green rows, with two consistent states: red columns over green rows, or vice versa.

**Figure 9:**
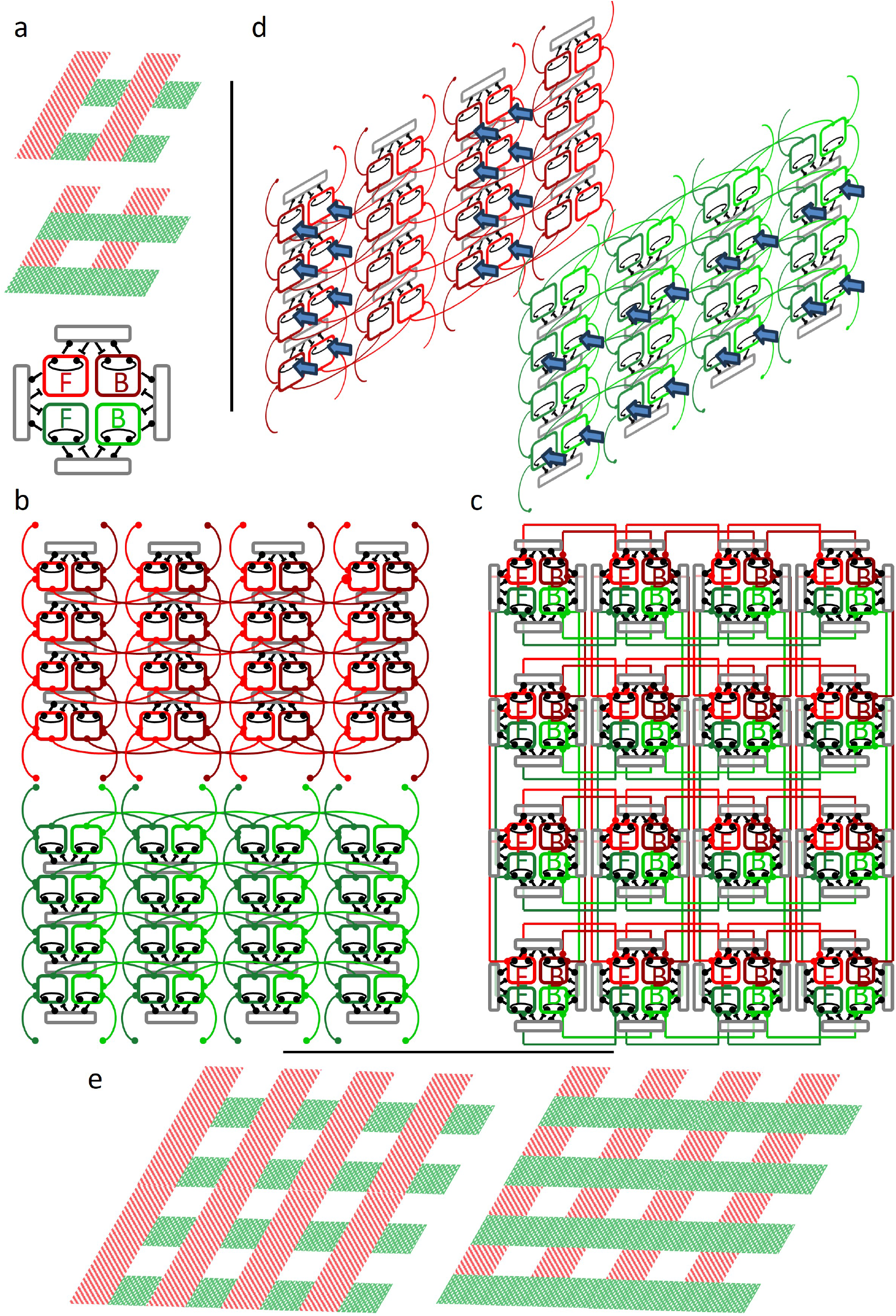
2-D laterally connected PCRNs for gratings bi-stability. **a.** Two possible interpretations of a 4 *×* 4 grating. There are 16 blocks with four *P* = 2 PCRNs each sharing excitatory subpopulations (bottom figure in the panel) for a total of 64 excitatory subpopulations and 64 inhibitory pools enforcing local XORs: red and green foreground subpopulations in a block can’t be simultaneously winning or simultaneously losing. Globally, lateral connections make the system prefer states where either all the red blocks are in the foreground and all the green blocks are in the background, or the other way around. **b-c**. The PCRN forcing a color to be either in the foreground or background is laterally connected to the same color PCRNs in the four neighboring blocks. PCRNs in the same column of the first and last rows are assumed to be neighbors as well as the PCRNs in the same row of the first and last columns. Panel b shows the connectivity of the 2 *×* 2 grating without circular connectivity. anel c shows the full connectivity. **d**. Equal external stimulation is given to all the red PCRNs in the odd columns and all the green PCRNs in the even rows. **e**. Two possible interpretations of an 4 *×* 4 grating contains four copies of the network in panel c. See an animation in Supp. Fig.10.

The dwelling in or around a fixed-point is an inherently different dynamic state than that of the flowing between fixed-points. Positing that momentary perceptions and thoughts correspond to a series of state attractors obviates the need for ‘readout’ of internal representations of experiential states (Rumelhart et al., 1987), except those relevant for subsequent action. Attractors offer a simple and natural resolution to the ‘homuncular conundrum’ implicit in a discourse dominated by feed-forward models. In other words, ‘readouts’ are not needed for comprehension, only needed for motor response. Furthermore, Figs. 4(c,f) show that an ‘attractor state’ can comprise simultaneous and coordinated activity in a compound network of multiple elementary PCRNs, and that a set of elementary PCRNs can give rise to other such compositions (Supp. Figs. 9 and 10). Attractors can be thought as an explicit representation of the stimulus (even external stimulus), and the flow need not to ‘culminate’ in something like a ‘grandmother cell’ for comprehension. Early cortex comprehension can address a host of unanswered questions about our capacity to quickly form internal representations of new stimuli like QR codes and unfamiliar words from natural languages with different phonemes.

**Figure 10:**
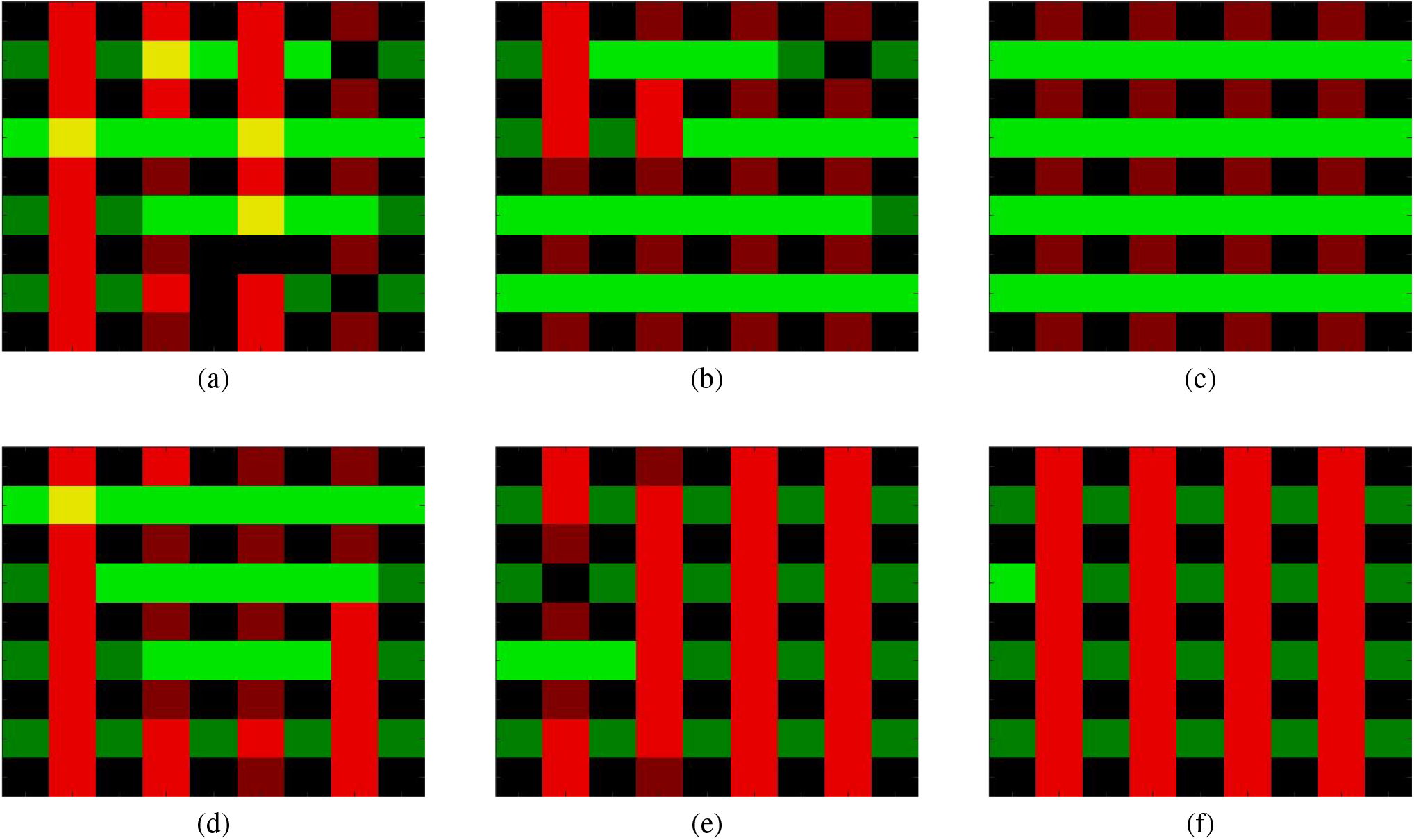
Animation - 2-D laterally connected PCRNs for gratings bi-stability. **a-f.** A 120 frame animation generated from a 6,0000 cycles simulation of a 4*×*4 grating compound PCRN. An extra-column and an extra-row of PCRNs have been added to the right and to the bottom of the grid depicted in Supp. Fig. 9. **a**. After 750 cycles. **b**. After 1,500 cycles. **b**. After 3,000 cycles. **c**. After 3,750 cycles. **d**. After 4,500 cycles. **e**. After 6,000 cycles. Animation can be found at this link.

Behavioral electrophysiology data often involves decision tasks with very similar options, leading to ambiguities. Our model addresses these scenarios and suggests that data may conceal sampling processes, even when they occur rapidly. Furthermore, the structure of ‘columns’ with their redundancy might play a significant role beyond cell death protection, potentially assisting in probabilistic sampling. This indicates that the Monte-Carlo method used in simulations also has practical applications in brain functions, allowing it to perform ‘Monte-Carlo’ simulations for various functions, configurations, or responses.

It is plausible to conceptualize cortical hyper-columns as PCRNs, a neural assembly encompassing a collection of competing attractors that share an inhibitory pool, representing the value of a specific local attribute derived from input stimuli (e.g., orientation). This notion suggests the hypothesis that hyper-columns within a particular cortical zone function as a network of laterally-connected PCRNs, consistent with physiological findings (Lund et al., 2003). The term ‘column’ in this context does not necessarily denote a section of cortex encompassing all layers. It is conceivable that the configuration of basic PCRNs is also confined to a single layer. For instance, the input layer (4) comprises sets of orientation PCRNs (e.g., purely monocular) that are distinct from those present in either more superficial or deeper layers.

## 6 Discussion

The explanatory power of the model has been illustrated here with findings from perceptual bi-stability, and its potential relevance for cognitive-level behavior is readily apparent. Until today, it has been a mystery what establishes when two percepts can coexist or when they suffer from XOR (from the behavioral point of view). Our model offers a computational principle which is also a prediction and requires verification: populations that encode different percepts and share an inhibitor pool are perceived in XOR. Conversely, precepts can coexist if the populations encoding them join separate inhibitor pools.

The model also offers new ways to think about the stochastic nature of transitions, and new tools to reexamine experimental discoveries that can shed light on the relationship between time-scales and physiological factors. The notion of ‘noise’ has been accumulating popularity (when compared to adaptation), but this model sets a new standard. Rather than introducing ad-hoc noise (as a supplement or appended), transitions happen because of the most basic and accepted model of noise: Poisson spiking of individual neurons and asynchronous updates. That’s all. This means time-scales are established through other factors, not noise. This new interpretation opens a series of predictions that seek verification regarding the physiological factors that must be influencing the transition times: input strengths as well as connection strengths between the different populations, and the structure of intra-cortical connectivity such as overlaps among excitatory neurons. The architecture is consistent with discoveries of cortex organization (Lund et al., 2003; Hubel and Wiesel, 1977). Feed-forward PCRNs are referring to inter-regional computations, and can explain the qualitative differences between the typological characteristics of inter-regional links - uniform, from those of intra-regional - intricately patterned. The computation within a region of a network of multi-PCRNs also explains the observations of Hubel and Wiesel (Hubel and Wiesel, 1977) regarding the typical characteristics of short-term versus long-term connections among hypercolumns.

The neural abundance and dense connectivity, necessary to create the balance (and probability through alternations) is indeed one of the earliest modern era observations about the brain (Ramón y Cajal, 1911), implying that evolution equipped the cortex of the mammals with the multiplicity needed for its functions. It is then, also natural to ask why evolution has converged to a PCRN architecture, given the seemingly simpler formulation of the symmetrized system. Beside the physiological constraints, specifically the finding that neurons’ pre-synaptic transmitters can be either excitatory or inhibitory, but not both [Dale’s Law], this model offers a more principled, computational justification, which stems from the need to learn and adapt to changing conditions in the external environment. Consider a system tuned to reflect that two stimuli, A and B, are equally likely. Now imagine that two types of changes occur simultaneously: one is a change in the prevalence of the two stimuli from [p(A),p(B)]=[0.5,0.5] to [0.4, 0.6], and the other is an increase in the overall magnitude of both stimuli, which would lead to an increase in the overall activity in the system. Physiological studies have found several global homeostatic processes geared towards keeping the overall level of activity in the system at a stable level, a goal which can be easily achieved in the PCRN by adjusting the value of *K*. At the same time, the change in the relative prevalence of stimuli *A* and *B* can be learned in an unsupervised manner with a simple Hebbian rule which incrementally changes the internal connectivity of the two populations, 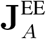 vs 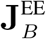 (Supp. Fig. 13 illustrates the feasibility of these parallel tracks of learning in a PCRN). In contrast, the values of *K* and **J**^EE^ are expressed by a single synapse per neuron in the symmetrized system, precluding the ability to address different environmental changes with separate learning mechanisms. Moreover, the numerical value of changes to the *K* and **J**^EE^ components of the synaptic weight can depart by orders of magnitude, causing a need to control the synaptic strengths at a high digital depth over a wide range of values (Supp. Fig. 13, panels (a) and (c)). The PCRN architecture thus offers significant advantages when learning and adaptation are taken into account.

**Figure 11:**
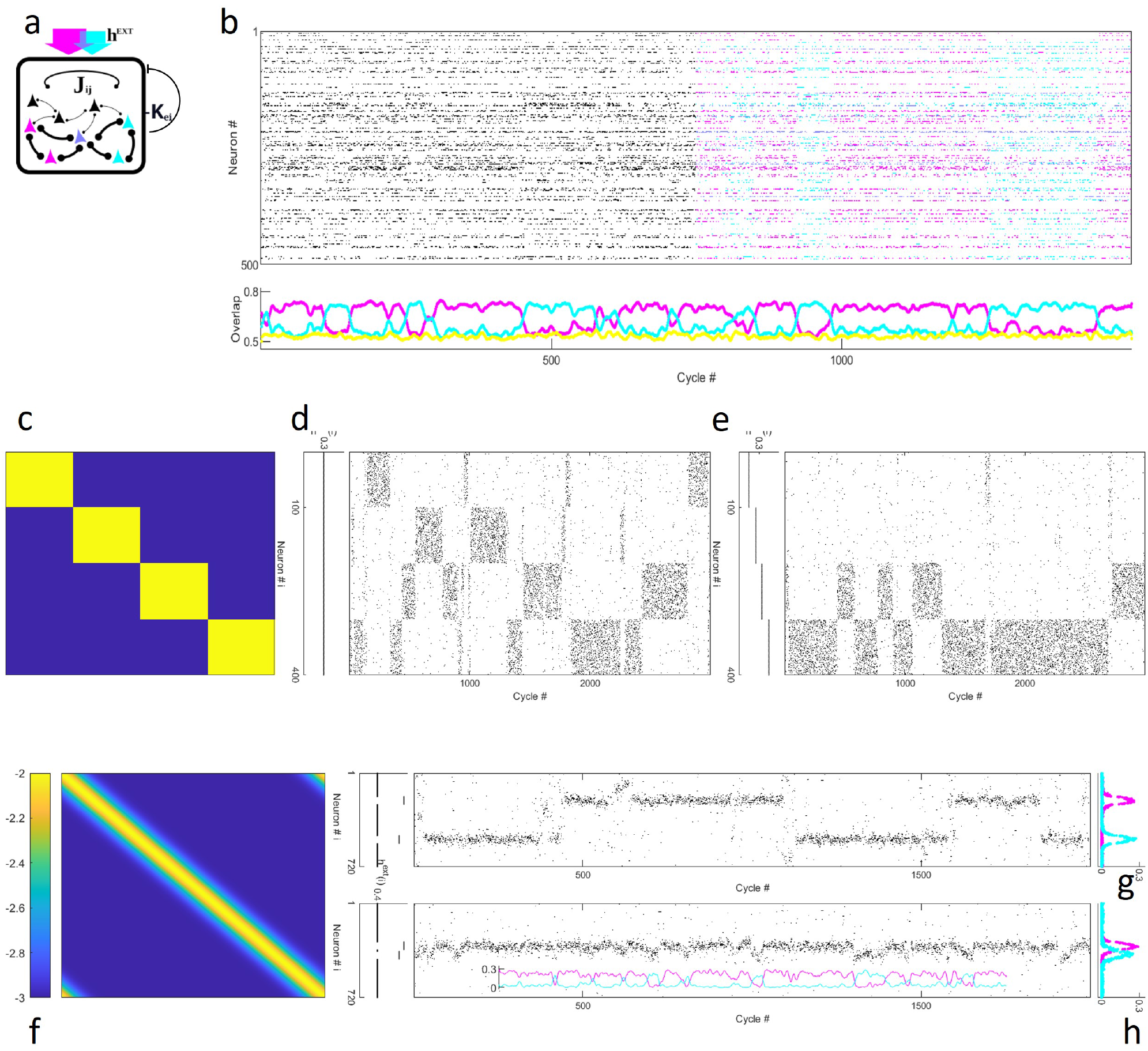
Simulation with symmetrized PCRNs. **a.** The symmetrized PCRN model architecture. An excitatory neural population is interconnected by the same type of Hebbian synapses than in the standard PCRN (positive connections), but to which a constant inhibitory value (*K*_ei_) is subtracted to their excitatory weight. Reciprocal inhibitory connections (negative connections) with the same constant inhibition value are also added between neurons across populations (*−K*_ei_). **b-h**. All simulation panels (b, d-e, g-h) from main Fig.1 are replicated using a symmetric connectivity (**J**^AF^*≡* **J**^EE^ *−K*_ei_J^N^, see main document Eq(2)). Parameters are not changed except for temperature that was increased by the same constant factor across all simulations. Because of the stochasticity of the asynchronous updates, the number of active inhibitory and excitatory neurons needed to maintain balance can get slightly misaligned. Nevertheless, and although the ‘regulatory’ task of the inhibitors might lag behind (see Supp. Fig. 12), if the temperature is not too high, the inhibitory population will catch up and will keep the activity of the excitatory neurons under control. This misalignment doesn’t occur in the symmetric system, dropping its contribution to the probability of alternations. Hence, if the temperature is not increased, there are no alternations. **b**. Ten random binary patterns were embedded in an excitatory population of 200 neurons, and 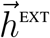 targeted the neurons belonging to two of the patterns, with unequal strength. Sampling behavior is hard to discern from the raster (top) when neurons are unlabeled (cycles 1-750, monochrome), but clear in the labeled portion (cycles 751-1500) and in time-courses of the overlap between network state and each pattern (bottom; yellow shows an unstimulated pattern). **c**,**f**. Connections within the excitatory population are symmetric (**J**^AF^). There is no inhibitory population. **c-e**. With four binary patterns of contiguous, non-overlapping sites, sampling, as in the non-symmetric case, appears in the rasters as intermittent blocks of activity and quiescence. Under uniform stimulation (panel d, flat black line), each of the competing sub-populations is active a quarter of the time, on average. When *h*^EXT^ values differ between populations, the network spends more time in those more strongly stimulated (panel e). **f-h**. Sixty densely overlapping Gaussian patterns in a ring, embedded in a symmetrized PCRN of 600 excitatory neurons, produce a smooth diagonal ridge in the connectivity matrix (f). Simulations show the same behavior as in the non-symmetric architecture. Stimulation at two well-separated loci results in visible alternations between distinct neuronal groups (g), but states remain differentiated even with minimally-separated inputs (h; magenta and cyan time-courses show mean activity in neuronal subsets stimulated by higher and lower peaks, respectively). In both cases, neurons’ activity profiles (right-most panel) do not mirror the boxcar shape of external stimulation (black lines, left), but rather follow the embedded patterns, indicating the same dynamics of the non-symmetric system governed by attractor states. Here, as Fig.1, *J*_IE_ =1, and *K*_EI_ =3. (For remaining parameters see Methods.)

**Figure 12:**
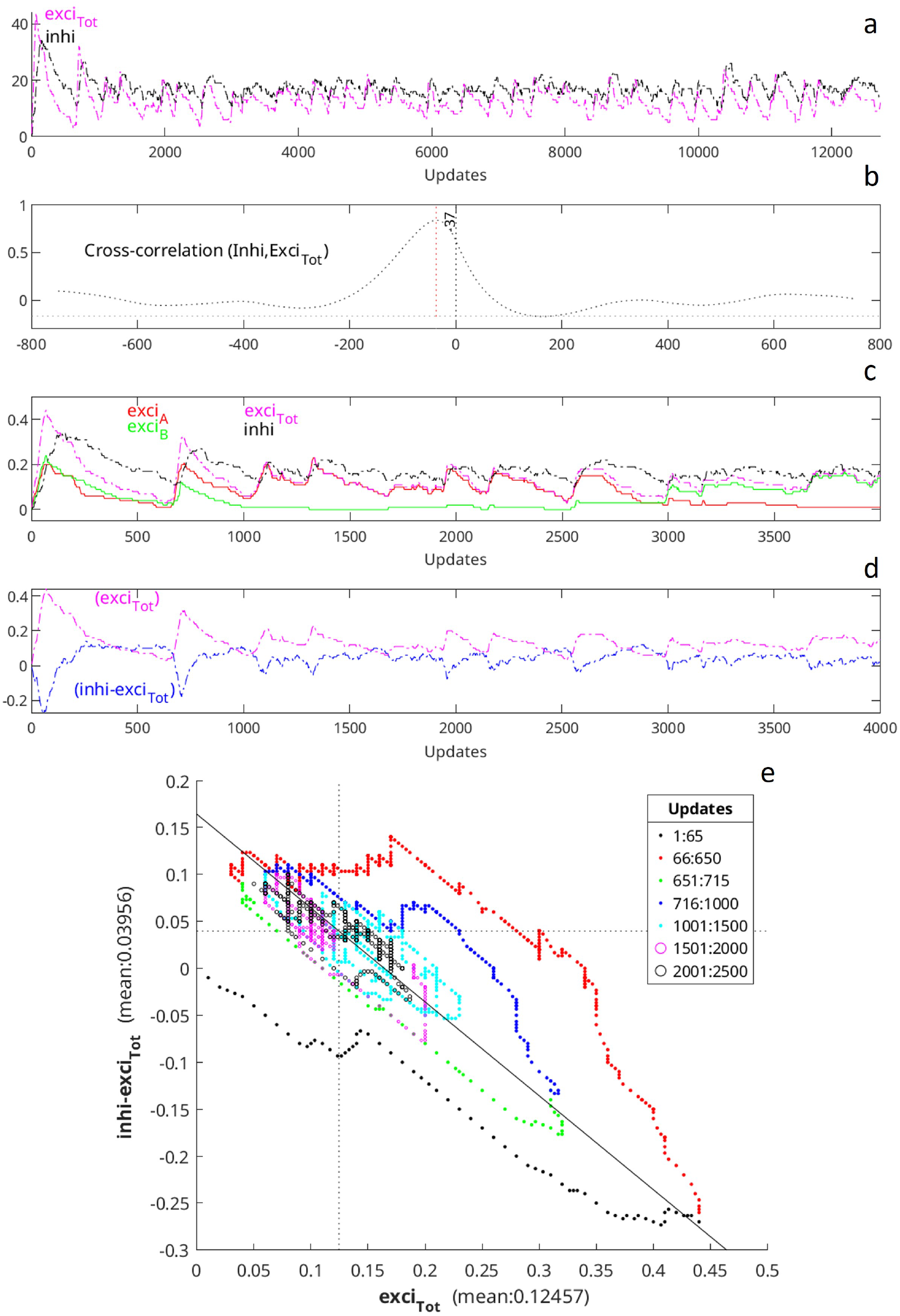
Inhibitory activity lags behind excitatory activity. **a-e.** Simulation of 200 cycles of a *P*=2 PCRN, with *N*_*EP*_ =*N*_*I*_=100, for a total population, *N*, of 300 neurons, *h*^ext^=0.3, and temperature *T* =2. Calculations for all panels are made based on the sequence of random selections of individual neurons considered for update. There are *N* of such checks in one cycle. The system starts with all neurons inactive (0). **a**. Time-32 courses plotting the total activity of the excitatory (exci) population (in magenta) and the activity of inhibitory (inhi) pool (in black) during the first 13,000 updates. They are clearly correlated with the inhi activity often above the exci, and it is consistent with the physiological observations reported in (Okun and Lampl, 2008). **b**. Normalized cross-correlation between the activity of the exci population and the inhi pool showing that changes in the inhi activity lag 37 steps behind the exci activity. **c**. Normalized activities (by *N*_*EP*_) separated by exci subpopulations over the first 4000 updates. Up to around step 1000, there are points where the exci activity is visibly higher than the inhi activity. After that, it happens less often and the difference is smaller. **d**. Time-course plotting the difference between the inhi and the exci activities (blue dotted line). (It would be 0 at temperature 0 in the balance regime after reaching a fixed-point). This difference starts with sharp fluctuations, but, as the system moves along the simulation, it quickly moves to values close to 0. The mean is marked with the dotted line at 0.04. **e**. Scatter plot of (normalized) exci activity against inhi minus exci activities. The simulation starts with an initial state of 0 activity, point (0, 0). During the first 65 steps (black dotted line), the external stimulation gradually activates exci neurons, and in turn the exci activity activates inhi neurons, but it happens at a slower pace (see panel c), hence, the y-axis values are negative. Around step 66 (orange dotted line), the inhi activity starts to affect the exci population and its activity starts to decline. The inhi activity continues to increase until it catches with the declining exci activity (panel c), at which point the inhi activity also starts to decline. Around step 650 (red dotted line), the exci activity starts to increase again but soon the *A* subpopulation pulls ahead and becomes the winner. Soon after, the inhi activity rises and peaks around step 780, just after the exci activity has peaked around step 715, and from there (blue dotted line) it decreases towards a ‘steady-state’ mean. At 0 temperature, the mean activity of the exci population at a fixed-point is 0.15. At higher temperatures, the mean activity is lower (in this simulation is 0.12 – see Supp. Fig. 5), and there are alternations between fixed-points (one is observed around step 3000).

**Figure 13:**
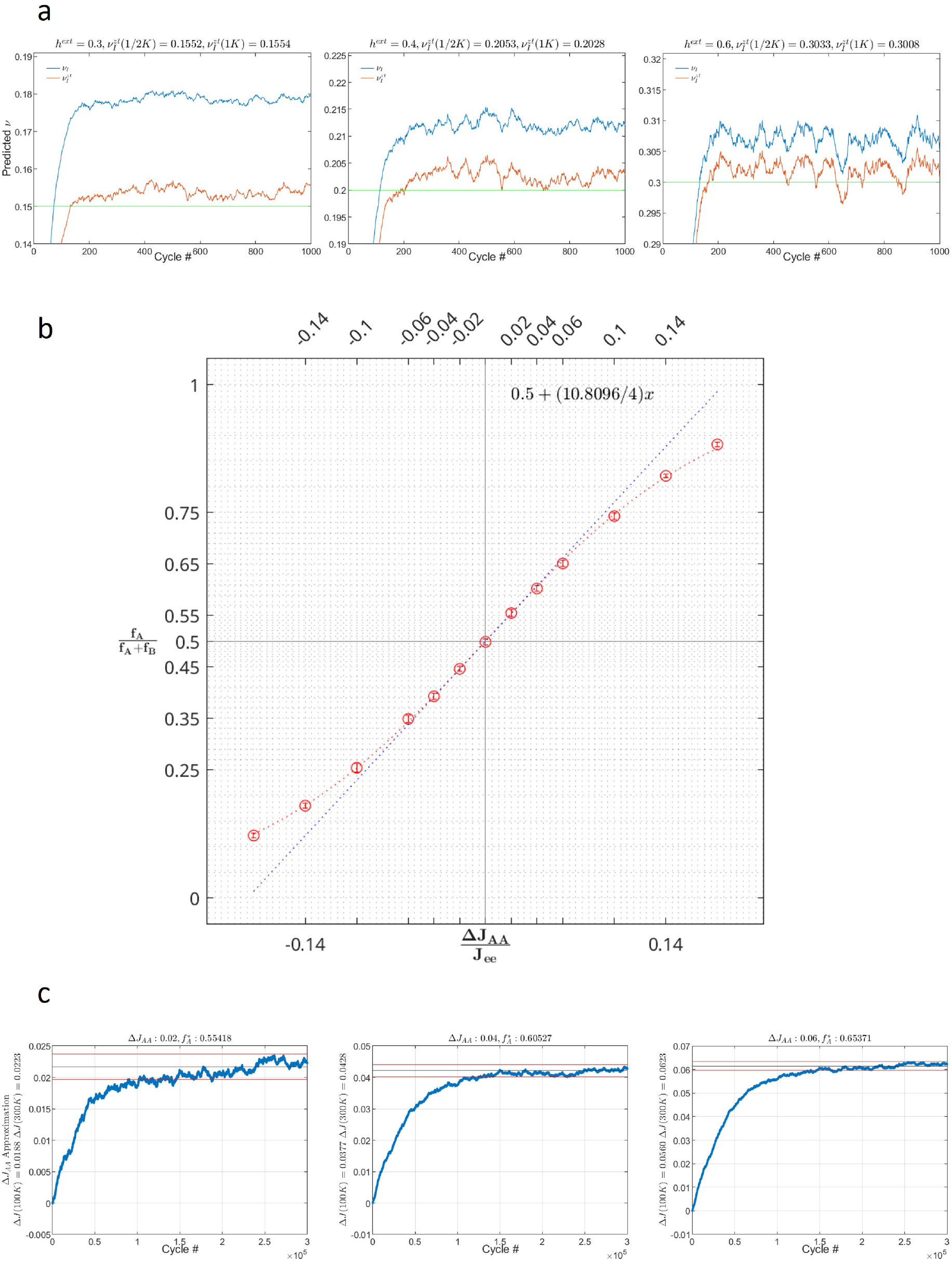
Learning J^EE^ vs K. Parameters *K* and **J**^EE^ have distinct roles in the dynamics of a PCRN. *K* controls overall exci population activity, acting as a normalization factor to changes in stimulation strength (eg. changes in contrast). Thus, *K* changes are driven by exci population activity changes within the PCRN. Conversely, **J**^EE^ is set to let the PCRN sample a probability distribution that reflects structural features of external stimulation. We illustrate the two cases using a simple *P* = 2 PCRN with *N*_*EP*_ = *N*_*I*_. To keep the mean firing rate of *E* around a fixed value, *ν, K* can be increased (or reduced) when the firing rate goes significantly above (or below) *ν*. Since the activity of *I*, the inhi pool, closely follows the activity of *E* (supp.Fig.7), the adjustment can be done based on changes of *ν*_*I*_. With the current *K* and the target mean activity *ν*, the new *K*^*new*^ can be approximated by 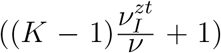 (see main paper, Eq. 7). For a fixed temperature 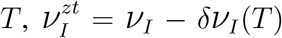. *ν*^*I*^ can be estimated by simply estimating the expected number of active neurons in *I* after an evaluation cycle. Denoting by *V*_*I*_(*t*) the percentage for such activity in cycle *t*, the Exponentially Moving Average formula, *ν*(*t*) = *ν*(*t −* 1)[1 − *α*] + *αV*_*I*_(*t*), with an arbitrary initial value *ν*(0) and *α <* 1, gets us *≈ ν*_*I*_, for *t* sufficiently large. Using the equations just above Eqs. 10 and 11 of Supp. Note 2, we get 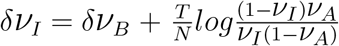. *V*_*A*_ and *V*_*B*_ refer to the mean activity of the winning and loosing subpopulations respectively. Since 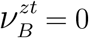, then *δν*_*B*_ = *ν*_*B*_, and both *V*_*A*_ and *V*_*B*_ can be estimated similarly to *ν*_*I*_. Furthermore, for fixed *T* and *h*^ext^, these values are quite stable, and a few hundreds cycles are sufficient to get good approximations. **a**. Simulations of 1000 cycles to estimate *V*_*I*_ and 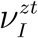 for three values of *h*^ext^, 0.3, 0.4, and 0.6, with *K* = 3, *T* = 30, *N*_*EP*_ =*N*_*I*_ = 500, and *α*= 0.025. Estimations are shown after 500 and 1,000 cycles. **b-c**. The behavior of a *P* = 2 PCRN is observed to learn **J**^EE^. That is, *f*_*A*_, the fraction of time of finding subpopulation *A* winning in a PCRN receiving uniform external input with **J**^EE^ as its exci connectivity matrix, will be (approximately) the same as *f*_*A*_ in the instigating PCRN. **J**^EE^ can be indirectly learned through learning the appropriated Δ*J*_*AA*_, and setting the *A* and *B* sub-network connectivity within **J**^EE^ to 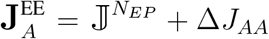, and 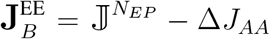. **b**. Simulation results showing the Sigmoid curve that captures the relationship between changes in Δ*J*_*AA*_ and *f*_*A*_ in a *P* = 2 PCRN with *N*_*EP*_ =*N*_*I*_ = 500, 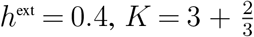, and *T* = 30. The panel also shows the linear approximation 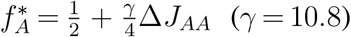. **c**. Simulation results to estimate Δ*J*_*AA*_ from three PCRNs with the same parameters used in panel b, having approximately *f*_*A*_ values 0.55, 0.60, and 0.65. Denoting by *V*_*A*_(*t*) (resp. *V*_*B*_(*t*)) the number of active neurons in *A* (resp. *B*) at cycle *t*, let *R*_*A*_(*t*) = 1 if *V*_*A*_(*t*) *> V*_*A*_(*t*), or 0 otherwise, and let *R*_*B*_(*t*) = 1 *− R*_*A*_(*t*). Note that 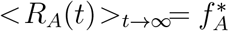. Taking *y*(*t*) = *c*(1 *− e*^*αt*^), *y*(0) = 0, and *y*(*t* + *dt*) = *y*(*t*)(1 *− α*) + *αc, y*(*t*)_*t→∞*_ = *c*. Replacing *c* with 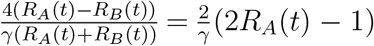, it can be shown that 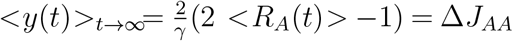. Results after 100K and 300K cycles with *α*= 2.5*×*10^*−*5^, *γ* = 10 show that significantly more cycles are needed to learn Δ*J*_*AA*_ than *K*, due to the many state transitions required for accuracy, unlike *ν*_*I*_, whose value remains stable after a few cycles if *T* and *h*^ext^ remain unchanged.

**Figure 14:**
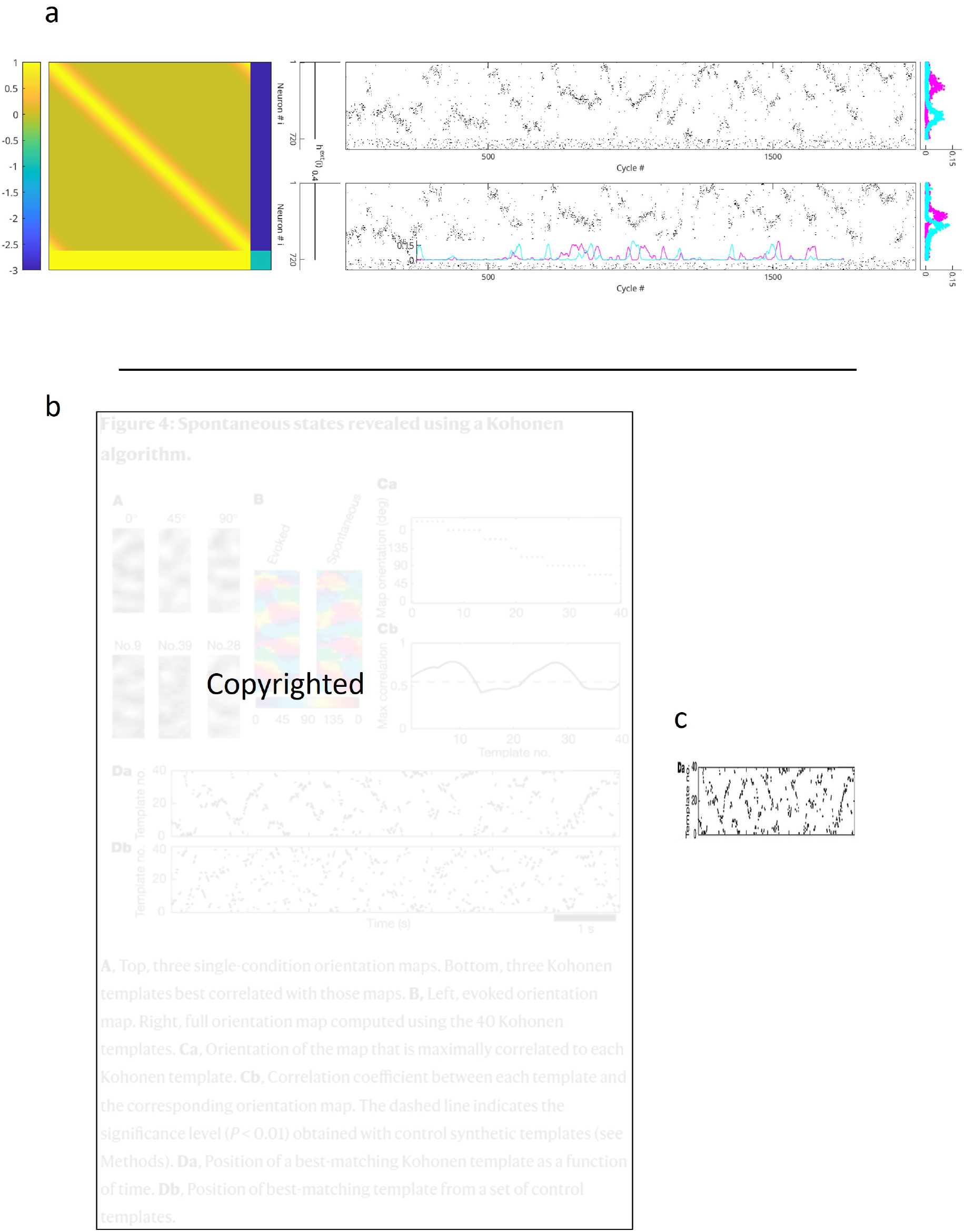
Spontaneous cortical activity and in the Gaussian ring PCRN. **a.** Simulation using a flat external stimulation on the PCRN embedding the sixty densely overlapping Gaussian patterns of Figs. 1f-h. The two rasters are for the same simulation, and it shows the spontaneous activity of neurons moving in waves. There are no visible alternations between distinct neural groups, but activity data was collected at the same two well-separated loci selected for Fig.1g and the two close loci selected for Fig.1h. The results plotted on the two graphs on the right show that the activity of the groups remain differentiated, recovering the shape of the patterns. The magenta and cyan time-courses in the bottom raster show mean activities in the two closely located neuronal subsets visualizing the segments where one of the them is winning. **b**. Reproduction of Figure 4 from (Kenet et al., 2003). Spontaneous occurrences of different states corresponding to cortical representations of orientations in cats were analyzed, states would correspond to occurrences of different patterns in a PCRN. The bottom rasters, Da and Db, in Fig.4 show how the spontaneous activity moves through the orientation templates designed by the authors resembling the wave patters in the raster of the PCRN simulation. **c**. Narrower version of Da from panel (b) to emphasize the waves.

**Figure 15:**
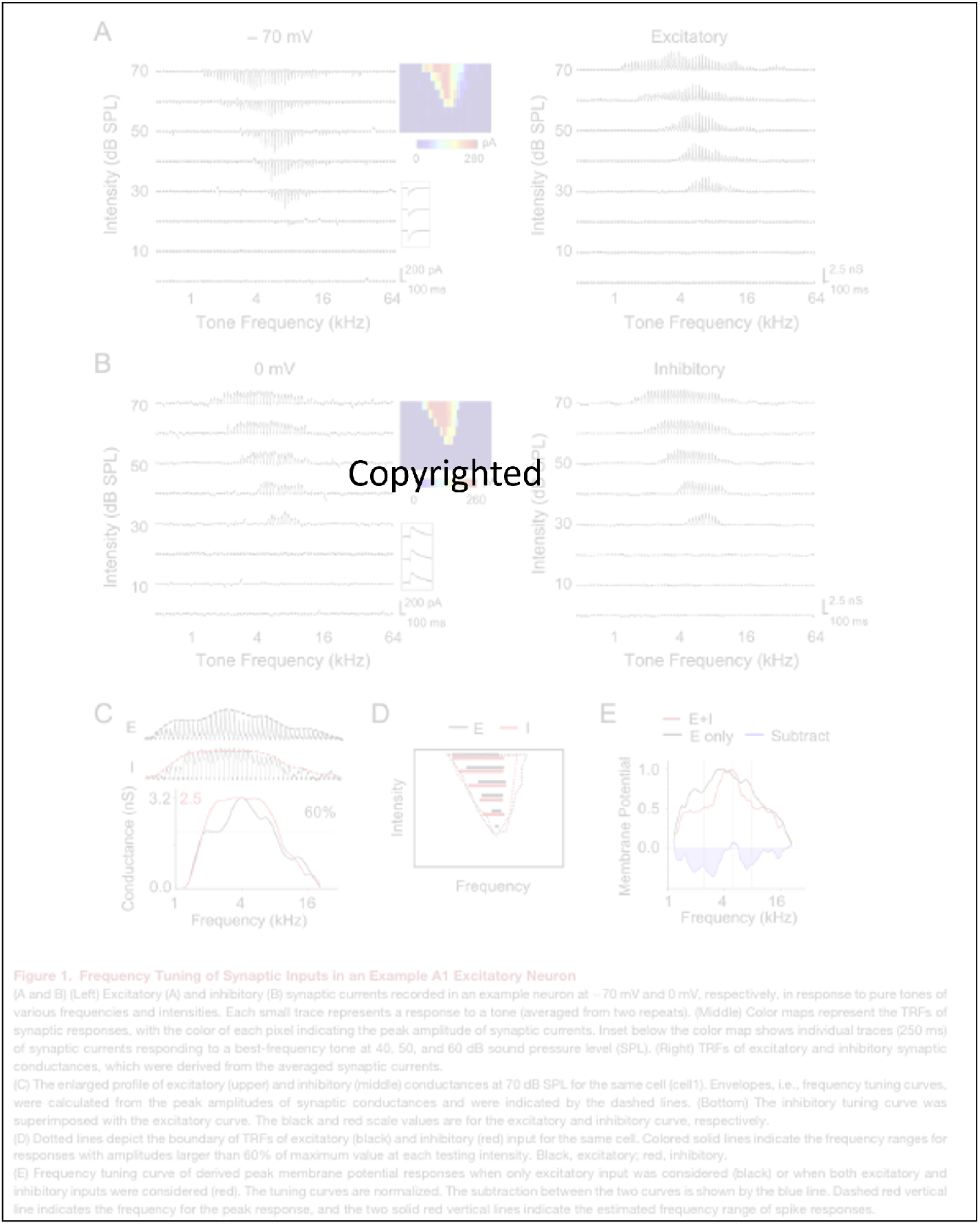
Physiological observations of Gaussian patterns. Reproduction of Fig. 1 from (Wu et al., 2008), Frequency Tuning of Synaptic Inputs in an Example A1 Excitatory Neuron.

The progression from single neurons to assemblies comprising hundreds of binary neurons may seem restricted by the limitations of current VLSI technology employed in artificial intelligence systems. However, recent advances in the field of nano-optics (Shen et al., 2017) offer promising compatibility with such architectural innovations.

## 7 Materials and Methods

Experimental verification of the theoretically predicted model dynamics used MATLAB simulations of neural networks with Eq. 3 as the update rule. The rule and the asynchronous neuron selection are implemented using the `mrg32k3a` random number generator. The PCRN parameter configurations used to generate the data in the figures are provided in the main text and figure captions. Reviewers can access the code via a Zenodo record as a direct ZIP download (no account required) at this link.

## Data availability

The code will be released under the AFL v3.0.

## Appendix

### Note 1: factors affecting the sampling rate

For better understanding of the model’s key ingredients, it is useful to examine the interplay between external stimulation, the strength of inhibition and neurons’ internal noise. The large *x − y* plane in Supp.Fig. 6 spans the parameters 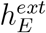 and temperature (= 1*/β*), respectively, for the case when the two populations receive external stimulations of equal strength. Each of the small tiles shows the solution(s) for the pair of parametric values corresponding to its location on the plane, with the remaining parameter held constant at *K*= 3. (The bottom row can thus be thought of as the low-temperature counterpart of the main diagonal in Fig.2(c); the solutions match closely, but note that here they were derived numerically.) This modified version of a phase diagram is informative about the gradual, quantitative changes in phase plane structure as well as the qualitative phase-transition lines. The * marks the region in parameter space used for the simulations of Fig.2(c). (The varying of Δ*h*^*ext*^ to obtain 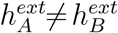 is thus equivalent to moving in the direction perpendicular to the *x− y* plane in Supp.Fig. 6.)

The effect of stimulation strength, on the other hand, may be less intuitively obvious. As 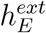 is increased, a slow-down of alternations is observed, in spite of there being no change in temperature. The reason is that stronger 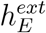 means higher *V*_*E*_, which reduces the overlap in activity levels between the two populations in turn reducing the probability of alternations. Importantly, the increase in *V*_*E*_ caused by stronger stimulation also entails a move away from the intermediate-*ν* regime, where inhibition from the common pool *I* counters the excitatory inputs in near-perfect balance. The transition to the ‘winner take all’ regime 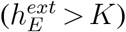 is reflected externally by *ν* hitting 1 (for the winning population) and the cessation of alternations, but the underlying reason is the inability of the inhibitory pool to keep up with mounting excitatory input. These two aspects of the model – probabilistic sampling, and the balancing of excitation by inhibitory feedback – are thus two sides of the same coin, inexorably connected. The model is consistent with physiological evidence that inhibition tracks excitation closely and near-instantaneously Okun and Lampl (2008).

The column of three rasters on the left also illustrates the effect of changing temperature on the dynamics. As may be expected intuitively, the alternations speed-up as the temperature, ie, neurons’ internal noise level, increases. For a stimulation value of 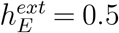, the alternations speed up as temperature is raised from 4 (bottom panel and large asterisk) to 5 and 6 (middle and top panels) until alternations between the two states start to be come difficult to separate. Nevertheless, as long as the parameters are within the shaded region, ie, the balance regime, the network’s ongoing activity conveys choice-relevant information.

Other rasters are shown in different locations within the region where the two attractors exist, but the essence is that there are three major engines driving variability: the balance kept between excitatory and inhibitory populations that can be broken if the external stimulation is too high, the sparking probability of each neuron in isolation, and ambiguous or conflicting input stimulating neuron subpopulations that can’t be active simultaneously. The three must be active, otherwise there are no alternations. As mentioned earlier, if the balance is broken *ν* will be 1. If the temperature is 0, the system will get trapped in one of its attractors, and variations of mean activity will be of order 1, caused by a ‘frustration’ mechanism. And finally, a non-ambiguous stimulus means that only neurons belonging to the same attractor are receiving input and all will have the same *ν* modulated by the strength of the input. Changes in dynamics are also affected by the intersection between excitatory subpopulations including ‘continuous’ connective of line attractors with different profiles, intra-connections between excitatory subpopulations, discretizations of external stimuli like in Figs.1(g-h) and Fig.3(j).

### Note 2: derivation of mean activities at low temperatures

Let’s assume a *P* = 2 PCRN with excitatory subpopulations, *A* and *B*, of size *N*_*EP*_ = *N*, and an inhibitory pool, *I*, of the same size. With no loss of generality, let’s assume the system at temperature 0 is at a stable fixed-point where subpopulation A is winning with its neurons having a mean activity of 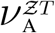. Therefore, the mean activity, 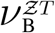, of neurons in subpopulation B is 0, and the mean activity, 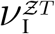, of the inhibitor population is 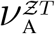.

Let 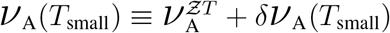, then

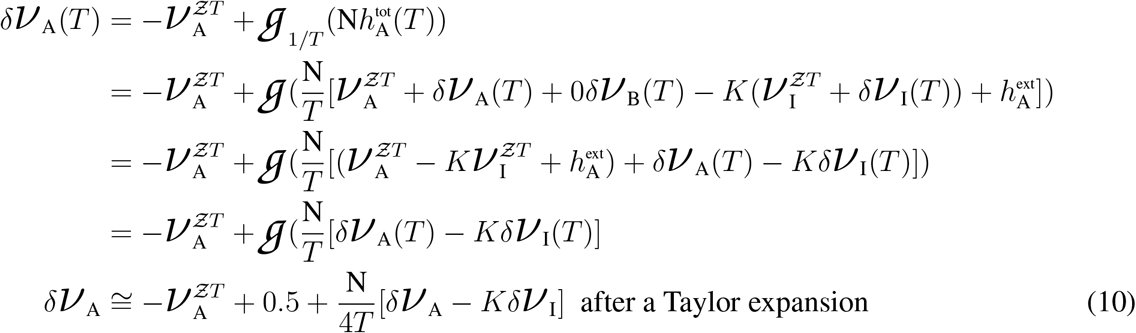

Let 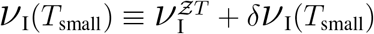, then

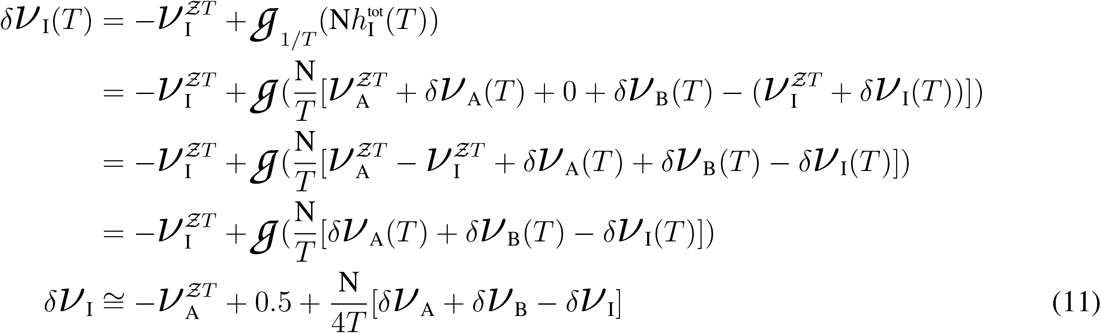

Let 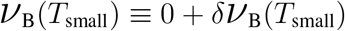, then

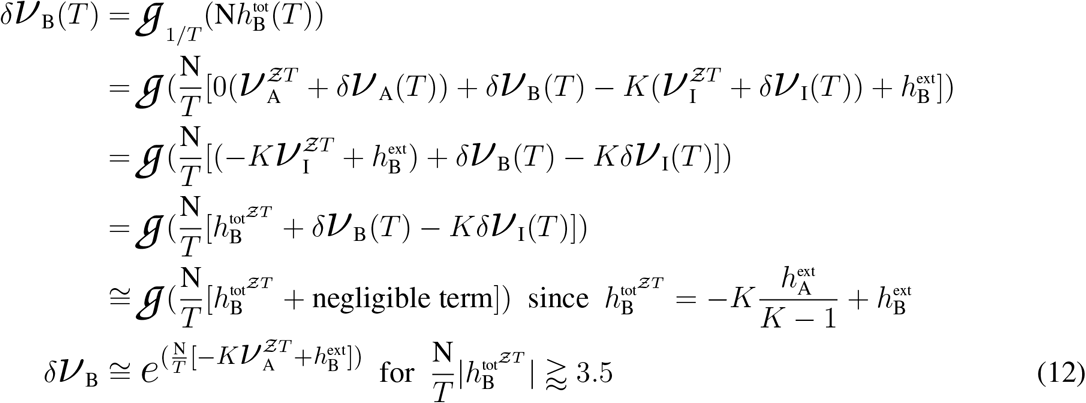

From Eqs.10 & 11,

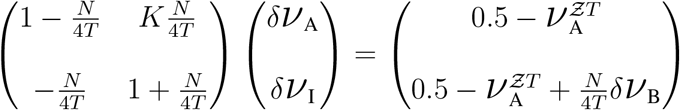

Letting 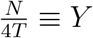,

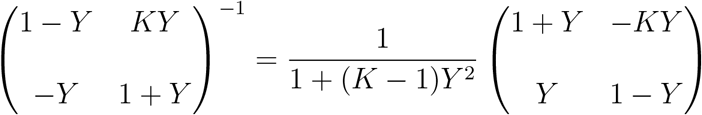

Hence,

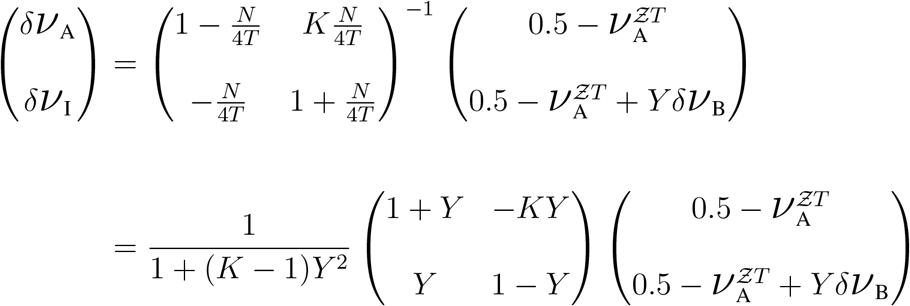

## Acknowledgments

Nava Rubin conducted part of this work as an ICREA Research Professor at Universitat Pompeu Fabra. Funding was provided by the Spanish Ministry of Economy and Competitiveness (TIN-2016-81032-P, MDM-2015-0502) and the European Union (PCIG11-GA-2012-322033). Jorge Lobo acknowledges the support of David Kleinfeld, Sebastian Seung, and Michail Tsodyks for reviewing the final manuscript.

## Author contributions

After N. Rubin’s death on November 20, 2024, J. Lobo used her notes and recordings to complete the Discussion section and the section on laterally connected PCRNs. He also implemented her detailed design notes for Figure 4 and Supplementary Figures 9, 10 and 13, developed several data-analysis utilities and visualization scripts, and wrote the legends for the supplementary figures.

## Notes

### Competing Interest Statement

The authors have declared no competing interest.

### Summary of Updates

The list of authors has been reordered to better reflect their individual contributions.

